# Neural microexons modulate arousal states via cAMP signalling in zebrafish

**DOI:** 10.1101/2025.04.24.650407

**Authors:** Tahnee Mackensen, Luis Pedro Iniguez, Thomas Soares Mullen, Cristina Rodríguez-Marin, François Kroll, Giulia Zuccarini, Jordi Fernandez-Albert, Laia Sancho-Vila, Jon Permanyer, Isaac H. Bianco, Michael Orger, Jason Rihel, Manuel Irimia

## Abstract

Arousal states, often dysregulated in neurodevelopmental disorders, shape how organisms perceive and respond to their environment. Here, we show that *srrm3*, a master regulator of neural microexons, is essential for normal arousal in zebrafish larvae. *srrm3* mutants exhibit persistent hyperarousal, including sleep loss, sensory hypersensitivity, anxiety-like behaviour, and heightened neural and behavioural activity. Elevated cAMP signalling likely drives this hyperarousal, as pharmacologically reducing cAMP rescues mutant behaviour, while increasing cAMP in wild-type larvae phenocopies the mutant hyperaroused state. Pharmacological cAMP modulation also mimics and reverses *srrm3*-dependent transcriptional changes. These include immediate early gene downregulation, which, together with altered activity-dependent transcription factor motif occupancy, suggest adaptation to sustained neuronal hyperactivity. Additionally, *srrm3* mutants show upregulation of microexon-containing genes, likely compensating for microexon loss. Together, these findings reveal a role for neural microexons in shaping arousal via cAMP signalling, providing insight into how splicing defects may underlie sensory and sleep disturbances.

## Introduction

Animals continuously adapt their behaviour in response to internal states and external environmental demands^1^. Two different behavioural states are sleep and wakefulness, with distinct baseline arousal levels. Arousal governs an individual’s readiness to interact with the environment by regulating central nervous system activation^2^. For example, high arousal may persistently increase an animal’s motor output in response to sensory stressors^3^, while low arousal (e.g. during sleep) decreases the threshold of responsiveness^4^. Sleep and wake states are conserved across the animal kingdom^5^, from jellyfish^6^, fruit fly^7,8^, and zebrafish^4,9,10^ to humans^11^, underscoring their evolutionary importance. Their broad conservation and emergence early in life^12,13^ suggest that arousal regulation relies on fundamental neural mechanisms specified during neurodevelopment. However, how these developmental programs arise and shape subsequent behaviour remains poorly understood.

Neurodevelopment is tightly regulated across time and space, both at the transcriptional level and through other molecular mechanisms such as alternative splicing. Alternative splicing allows for the generation of multiple transcripts from a single gene, thus greatly expanding protein diversity and function^14,15^. A particularly remarkable case of an alternative splicing program is that of neural-specific microexons^16,17^. Microexons, defined here as exons with 3 to 51 nucleotides (nts), are highly conserved across vertebrates^18^. Their splicing is controlled by the master regulators SRRM3 and SRRM4 through their enhancer of microexons (eMIC) domain located in their C-terminal region^16,19–21^. Microexons are included in a switch-like manner during late neurogenesis, fine-tuning processes such as axon guidance and synapse formation^16,22^. Therefore, they have the potential to shape neural circuits^23^, laying the foundation for balanced excitation and inhibition in the brain^20^. Microexon misregulation has been observed in neurological disorders such as autism spectrum disorders^16,24,25^ and schizophrenia^26^. Patients with these disorders commonly display difficulties in sensory processing^27,28^ and abnormal sleep-wake cycling^29,30^. Emerging evidence suggests that mis-splicing of neural microexons may directly influence sleep and arousal states. For instance, deleting a single microexon in the synaptic organizer *Ptprd* in mice results in sleep deficits and hyperactivity^31^. In zebrafish, double mutants for both *srrm3* and *srrm4* move more than their wild-type (WT) siblings^32^, and *Drosophila melanogaster* mutants lacking the eMIC domain of *Srrm234* sleep less, particularly at night onset^18^.

These findings suggest that microexons play important functions during neurodevelopment that affect the arousal system. However, it is unknown how widespread microexon mis-splicing gives rise to a coordinated shift in arousal-related behaviours, neuronal dynamics, and underlying neuromodulatory and transcriptional programs. To address these gaps of knowledge in a vertebrate system, we used a zebrafish homozygous mutant for the microexon master regulator *srrm3*. By 5 days post fertilization (dpf), zebrafish larvae exhibit a wide array of sensorimotor behaviours and a neurotransmitter system that fundamentally resembles that of mammals, including key arousal systems such as dopamine and noradrenaline^33–36^. At this stage, neurogenesis is still ongoing^37^, and microexons continue to shape critical developmental processes^16^. Here, taking advantage of the high-throughput and high-resolution behavioural tracking technologies available for zebrafish larvae, we observed that *srrm3* mutants show reduced sleep and hyperactivity, including sensory hypersensitivity and an overrepresentation of stress and anxiety related swim types, which together are consistent with a heightened baseline arousal state. These behavioural abnormalities were accompanied by an altered internal brain state, with increased baseline and stimulus-evoked neural activity. Using pharmacological interventions, transcriptomics and chromatin accessibility assays, we demonstrated that the heightened arousal levels upon microexon mis-splicing in *srrm3* mutants are tightly associated with increased cAMP signalling and transcriptional adaptation to persistent neuronal hyperactivity.

## Results

### *srrm3*^ΔeMIC^ larvae show hyperactivity, sleep loss, excess thigmotaxis behaviour and sensory hypersensitivity

To investigate how microexon misregulation during neurodevelopment impacts arousal states, we performed high-throughput behavioural tracking of 5 - 8 dpf *srrm3*^crg3/crg3^ zebrafish larvae that lack the *srrm3* eMIC domain^38^, which is necessary and sufficient for microexon inclusion^19,21^. This line was used throughout the study and the homozygous mutant is hereafter referred to as *srrm3*^ΔeMIC^, while the heterozygous mutant is referred to as *srrm3*^+/ΔeMIC^.

Measuring activity for *srrm3*^ΔeMIC^ across multiple days and nights, we observed a hyperactivity phenotype, with 61% more activity during the day and 66% more activity at night compared to WT siblings (Fig. 1A-C), which was also observed in a second founder line (*srrm3*^crg5/crg5^) (Fig. S1A,B). This hyperactivity, most prominent in the morning, was due to more frequent and longer swim periods, but not increased swim vigour, when compared to WT siblings (Fig. 1A,B; Fig. S1B). *srrm3*^ΔeMIC^ larvae also showed significant sleep disturbances, including a 47% delay in sleep onset (5.7 ± 1 min, model coefficient ± standard error [SE] from linear mixed effects model [LMM], p = 7e-5) and shorter sleep episodes (-0.4 ± 0.1 min, coefficient ± SE from LMM, p = 1e-5), resulting in an average loss of up to 1.5 hours of sleep at night compared to WT larvae (Fig. 1A,C; Fig. S1A,B). For some parameters, these effects were more pronounced in *srrm3*^ΔeMIC^*;srrm4*^-/-^ double mutants, while *srrm3*^+/ΔeMIC^*;srrm4*^-/-^ larvae did not show signs of hyperactivity (Fig. S1A,B).

**Figure 1:**
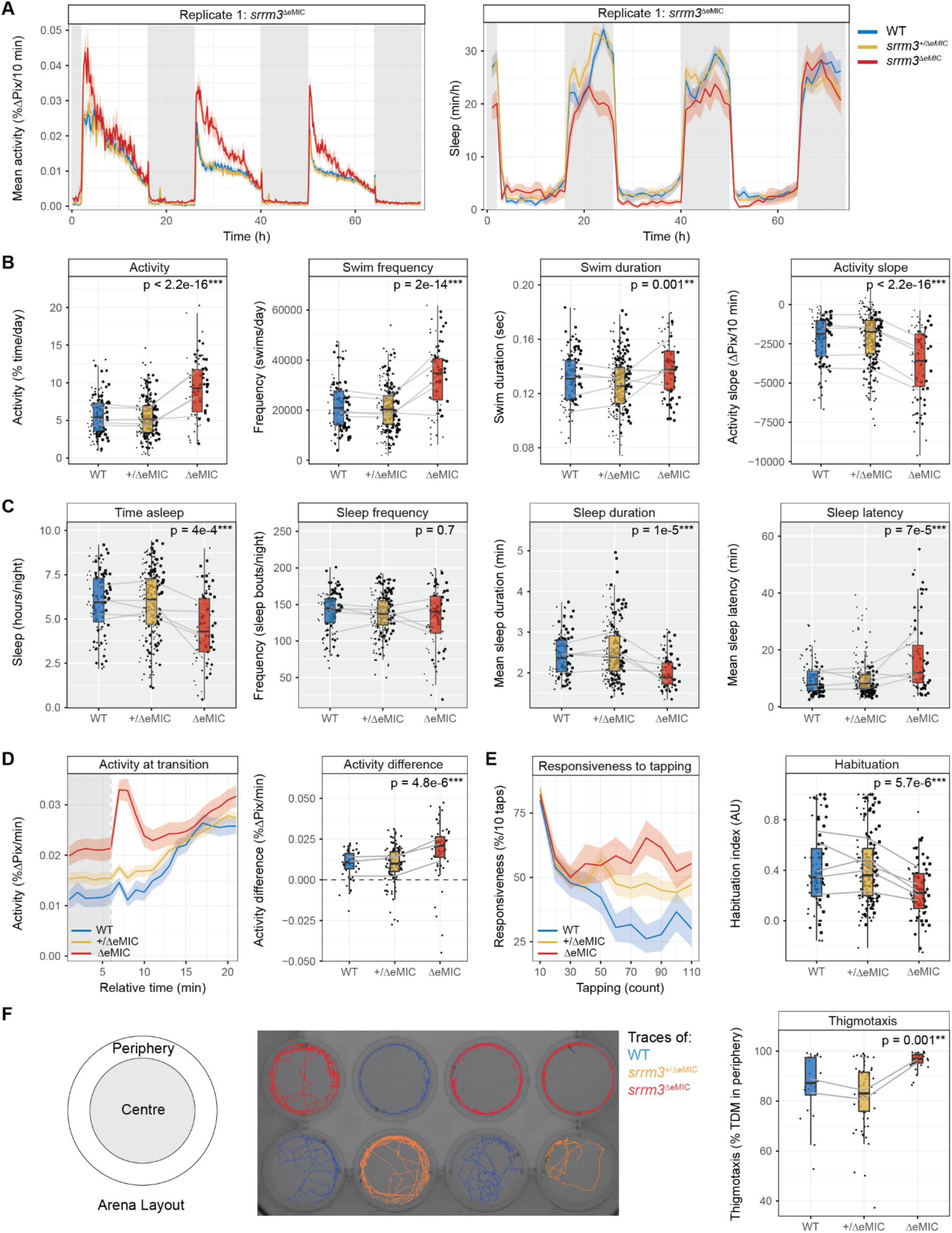
***srrm3*^ΔeMIC^ larvae show hyperactivity, sleep loss, sensory hypersensitivity and excess thigmotaxis. A:** Representative traces (mean ± SEM) for activity (%Δpixels [%ΔPix]/10 min) (left) and sleep (min/h) (right) during 74 h (5 - 8 dpf) on a 14 h light - 10 h dark cycle (white background and grey background respectively). For visualisation, values of refill periods were replaced with pre refill values. **B,C:** Behavioural parameters of activity (day [6 dpf]) and sleep (night [5 - 6 dpf]). Outliers were removed for visualisation only. N = 5 clutches (dot shape) with 13 to 57 larvae each per genotype. Statistics by likelihood-ratio test on LMM for day 6 dpf and for nights 5 - 6 dpf and 6 - 7 dpf^124^. **B:** Day: percentage activity: +3.5 ± 0.4% active/day, p < 2.2e-16; swim count: +11,866 ± 1,487 swims/day, p = 2e-14; active swim duration: +0.009 ± 0.003 sec, p = 0.001; activity slope: -1,562 ± 176 ΔPix/10 min, p < 2.2e-16. **C:** Night: time asleep: -0.8 ± 0.2 h/night, p = 4e-4; sleep frequency: 1.2 ± 3.4 sleep bouts/night, p = 0.7; sleep duration: -0.4 ± 0.1 min, p = 1e-5; sleep latency: 5.7 ± 1 min, p = 7e-5. **D:** Averaged (5 transitions per larvae) light-on response at 6 dpf. Left: Representative trace (mean ± SEM) of activity (%ΔPix/min) across larvae from the same clutch and genotype. Right: Activity (%ΔPix/min) difference (+0.010 ± 0.002 %ΔPix, p = 4.8e-6, LMM) 1 min after stimulus minus 1 min before stimulus. Sample size: N = 3 clutches (dot shape), with 12 to 47 larvae each per genotype. **E:** Habituation to mechanical tapping stimuli at 6 dpf. Left: Representative trace (mean ± SEM) of mean responsiveness (%) to 10 tapping stimuli at 90 sec inter-stimulus interval (tap 1-10) and 5 sec inter-stimulus interval (11 - 100). Right: Habituation index, where 1 indicates strong habituation (no response to the last 40 taps) and 0 no habituation (habituation index: -0.19 ± 0.04 AU, p = 5.7e-6, LMM). N = 5 clutches (dot shape) with 5 to 46 larvae each per genotype. **F:** Thigmotaxis assay at 6 dpf. Left: Arena schematic, divided into two equally sized areas (centre and periphery). Centre: Representative tracks (120 sec of movement) around the 40^th^ min of tracking. Right: Mean thigmotaxis (% TDM in periphery) during 1 h of tracking (+10 ± 3 % TDM in periphery, p = 0.001, LMM). N = 2 clutches with 10 to 43 larvae each per genotype. **B-F:** Statistics represent model coefficients ± SE with p-values shown for WT vs *srrm3*^ΔeMIC^ siblings. Each dot represents one larva. Lines connect genotype means across clutches. Detailed statistics and sample sizes in Table S1.

Although *srrm3*^ΔeMIC^ larvae responded strongly to light transitions (Fig. 1A,B), they were reported to exhibit severe vision deficits^38^. To test whether reduced vision can cause the hyperactivity phenotype, we compared locomotor activity in *vsx1*^-/-^;*vsx2*^-/-^ larvae, which have severe visual impairments, against *vsx2*^-/-^ siblings without visual abnormalities^39^. Both genotypes showed high sleep levels, and *vsx1*^-/-^;*vsx2*^-/-^ larvae exhibited reduced night-time activity relative to *vsx2*^-/-^ controls (Fig. S1A,B). This suggests that visual impairment alone does not result in hyperactivity or sleep loss, in line with multiple studies reporting hypoactivity in visually impaired and blind fish^40–42^ and indicates that the sleep/wake phenotypes of the *srrm3*^ΔeMIC^ larvae are not caused by their vision deficits.

Hyperactive zebrafish larvae, but also human patients with behavioural hyperactivity and sleep deficits, frequently present a greater stimulus-induced sensory response and heightened levels of anxiety^29,34,43–45^. We thus tested sensory responsiveness of the *srrm3*^ΔeMIC^ larvae to dark-light transitions and their habituation to mechanical tapping stimuli. In response to a sudden light-on stimulus, *srrm3*^ΔeMIC^ larvae displayed an exaggerated response, even considering their higher levels of baseline activity (Fig. 1D; Fig. S1C), and their reduced visual function^38^. Moreover, *srrm3*^ΔeMIC^ larvae showed reduced habituation to repeated mechanical tapping stimuli (Fig. 1E; Fig. S1D; -0.19 ± 0.04 habituation index (HI), coefficient ± SE from LMM, p = 5.7e-6). Finally, while thigmotaxis, or “wall-hugging” behaviour, a common proxy for anxiety^46^, was high across genotypes, levels in the *srrm3*^ΔeMIC^ were significantly above those of WT siblings (Fig. 1F; +10 ± 3% total distance moved (TDM) in periphery, coefficient ± SE from LMM). Together, these findings demonstrate that *srrm3*^ΔeMIC^ larvae exhibit a range of behavioural abnormalities, including hyperactivity, insomnia, anxiety-like behaviour, and sensory hypersensitivity, suggestive of a heightened arousal state.

### Movement repertoire and kinematics reflect a state of heightened arousal in *srrm3*^ΔeMIC^ larvae

To better understand the nature of the behavioural abnormalities in *srrm3*^ΔeMIC^ larvae, we examined their movement repertoire and kinematic features across sequences of swims, as these reflect the zebrafish larva internal state^47–49^ and can offer additional insights into hyperactivity phenotypes^50^.

Locomotion in larval zebrafish consists of discrete bouts of tail movements that can be categorized into a set of kinematically distinct types^50^. Tracking the tail movements of freely swimming individual larvae at high speed (700 Hz) under constant illumination (baseline) and, for some cohorts, during light/dark transitions (Fig. 2A), we identified 11 out of 13 possible bout categories, based on 73 kinematic parameters per bout^50^. The two categories not observed were capture swims only executed in the presence of prey^50,51^. The remaining categories can be grouped into: (a) low displacement forward swims, including approach swims (AS), Slow 1, and Slow 2; (b) low displacement reorienting swims, such as routine turns (RT), J-turns, and high-angle turns (HAT); (c) high-displacement swims, namely burst swims (BS), spot avoidance turns (SAT), O-bends, and short- and long-latency C-starts (SLC, LLC)^52^ (Fig. 2A,B; Fig. S2A).

**Figure 2:**
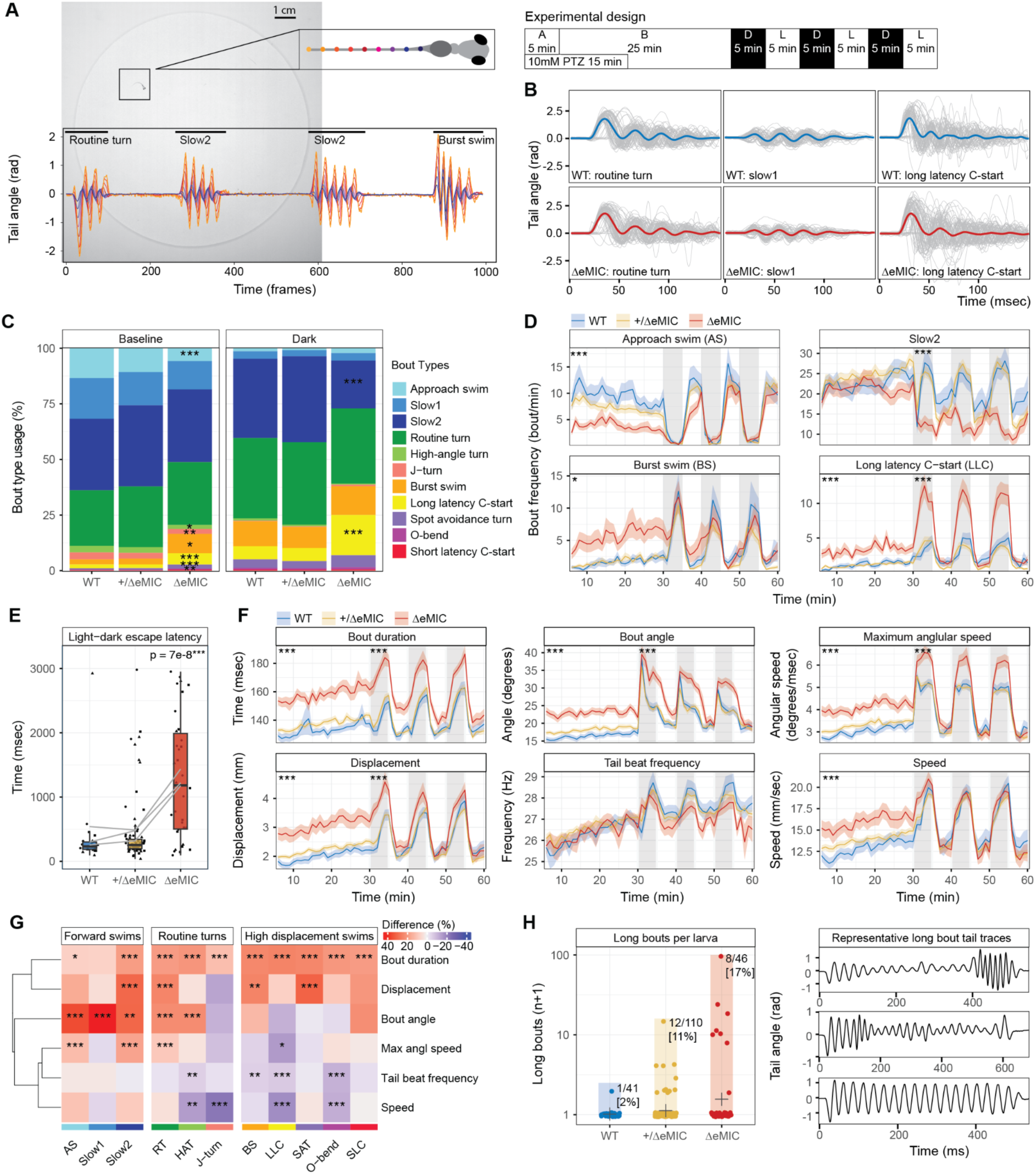
Bout type usage and kinematics reflects a heightened arousal state in *srrm3*^ΔeMIC^ larvae. A: Left: Circular arena with a 5 dpf larva. Larva diagram showing the 9 tracked tail segments (purple [anterior, 1] to yellow [posterior, 9]) each spaced 300 μm apart. Below, a representative WT bout sequence colored by tail segment. Right: Experimental design. Larvae from four clutches were tracked for 5 min (A = acclimatisation) followed by 25 min of tracking under constant illumination (B = baseline). Three clutches were further exposed to alternating 5 min intervals of dark (D) and light (L). Alternatively, larvae from two clutches were also tracked under the exposure of 10mM PTZ for 15 min. **B:** Representative tail traces (7^th^ tail segment) across 3 randomly chosen bout types with 70 randomly sampled individual traces per type (grey) and the mean trace shown in bold, from one clutch, with 8 WT and 14 *srrm3*^ΔeMIC^ larvae. **C:** Bout type usage (%) at 25 min baseline and 15 min dark. **D:** Mean bout frequency (bout/min) over time. **E:** Mean latency of first escape swim (O-bend, SAT, SLC, LLC) within the first 3000 ms after light-off (+235 ± 71%, coefficient ± SE from LMM, p = 7e-8). Dots represent larvae shaped by clutch. **F:** Mean bout kinematics over time, averaged over 1 min intervals. **D,F:** Grey background indicates dark periods. Traces represent mean ± SEM across larvae for each genotype. Significant stars are added for comparisons during baseline and darkness. **G:** Difference (%) in bout kinematic means within bout types. Values higher in *srrm3*^ΔeMIC^ compared to WT are displayed in red and lower values in blue. Statistics by maximum likelihood test on LMM. P-values were Bonferroni corrected for multiple bout types tested. N = 4 clutches with 3 to 33 larvae per genotype and bout type. **H:** Left: Long bouts (> 500 ms) by genotype. Numbers indicate the percentage of larvae with at least 1 long bout. Each dot represents one larva. The cross indicates the genotype mean. The y-axis is log-scaled and a pseudocount of + 1 was added for visualisation. N = 4 clutches with 7 to 34 larvae each per genotype. Right: Representative traces of long bouts. **C-G:** Statistics between WT and *srrm3*^ΔeMIC^ siblings by maximum likelihood test on LMM of log-transformed values **(C-E)** or untransformed values **(F,G)**. N baseline = 4 clutches, dark = 3 clutches, with 7 to 33 larvae per clutch and genotype. **C,D,F,G:** P-values in panels were Bonferroni-corrected for multiple testing. Significance stars represent: *p < 0.05, **p < 0.01, *** p < 0.001. Detailed statistics and exact sample sizes in Table S2.

Bouts in all 11 categories were observed in both WT and *srrm3*^ΔeMIC^ larvae, albeit at different frequencies (Fig. 2C). However, within each category, the mean and distribution for each bout type in a principle component (PC) space of bout kinematics^50^, was similar between the WT and *srrm3*^ΔeMIC^ individuals (Fig. S2B,C), unlike for larvae treated with the seizure-inducing compound pentylenetetrazol (PTZ) (Fig. S2C). This suggests no gross motor deficit in the *srrm3*^ΔeMIC^ larvae. We thus tested for a change in the choice of behaviour. Under baseline conditions, WT larvae rarely use high-displacement swims, typically associated with states of heightened arousal, for example during acute stress (Fig. 2C,D). In contrast, these swim types were drastically overrepresented among bouts of *srrm3*^ΔeMIC^ sibling larvae (Fig. 2C,D; Fig. S2D,E). When exposed to sudden darkness, a stress/anxiety-inducing condition, both genotypes increased the proportion of high-displacement swims. However, *srrm3*^ΔeMIC^ larvae continued to use high-displacement swim types at significantly higher rates than WT siblings (Fig. 2C,D; Fig. S2D; LLC: +160 ± 56%, percentage ± percentage SE from transformed LMM coefficients, p.adj = 1.7e-4). This observation is particularly striking since the *srrm3*^ΔeMIC^ larvae also exhibited a significantly increased escape latency to dark stimuli (Fig. 2E; latency: +235 ± 71%, percentage ± percentage SE from transformed LMM coefficients, p = 7.1e-8), suggesting heightened usage of high displacement bouts despite lower sensory responsiveness, a sign of hypersensitivity and increased baseline arousal^3,53^.

We next assessed whether *srrm3*^ΔeMIC^ larvae exhibit seizure-like swims, which would further reflect a shifted internal brain state in the *srrm3*^ΔeMIC^. For this purpose, we selected six kinematic parameters previously associated with seizure-like swimming patterns^54,55^ (Fig. 2F). Of these six parameters, all except tail beat frequency were increased in *srrm3*^ΔeMIC^ larvae compared to WT control siblings throughout the experiment. Particularly, bout duration was increased across all bout types (Fig. 2F,G; Fig. S2F). Interestingly, a subset of abnormally long bouts (> 500 ms) was observed in a significantly higher proportion of *srrm3*^ΔeMIC^ (17%; 8/46 larvae) than in WT larvae (2%; 1/41 larvae) (Fig. 2H; p = 0.03, two-sided Fisher’s exact test). Moreover, among the larvae that exhibited these bouts, *srrm3*^ΔeMIC^ ones executed them at a much higher frequency (Fig. 2H; ∼21-fold increase, p = 7.1e-08, two-sided Fisher’s exact test). These long bouts also showed a significantly larger bout angle, greater maximum angular speed as well as higher displacement, swim speed and tail beat frequency when compared to other bouts recorded for *srrm3*^ΔeMIC^ (p ≤ 0.001, permutation tests [see Methods]), thus presenting typical seizure-like characteristics^54,55^. Altogether, although *srrm3*^ΔeMIC^ larvae maintain a shared bout repertoire with WT larvae, they exhibit a profound shift towards stress-associated swims and an increased occurrence of rare, prolonged movements with atypical seizure-like kinematics, reflecting a heightened state of arousal.

### *srrm3*^ΔeMIC^ forebrain neurons are more active despite lower visually-evoked activity

Shifts in arousal states are reflected at the level of neural activity. This is most evident between sleep and wake^56,57^, but also apparent during the day, for example in response to visual stimuli^58,59^. Given the hyperactivity and sensory hypersensitivity of *srrm3*^ΔeMIC^ larvae, we assessed neural activity in different brain regions using two-photon (2P) calcium imaging, under baseline (no stimulus, grey screen) and visual stimulus conditions (light, dark, looming dot [LD], looming dot control [LDC]) (Fig. 3A). Notably, *srrm3*^ΔeMIC^ larvae with their heads restrained under the microscope remained hyperactive, with longer bouts across stimuli (Fig. 3B; +0.26 ± 0.04 sec, coefficient ± SE, generalized linear model [GLM], p = 1.8e-7) and increased bout frequency (Fig. S3A; +1.3 ± 0.6 bouts/epoch, coefficient ± SE from GLM, p = 0.02) compared to WT individuals.

**Figure 3:**
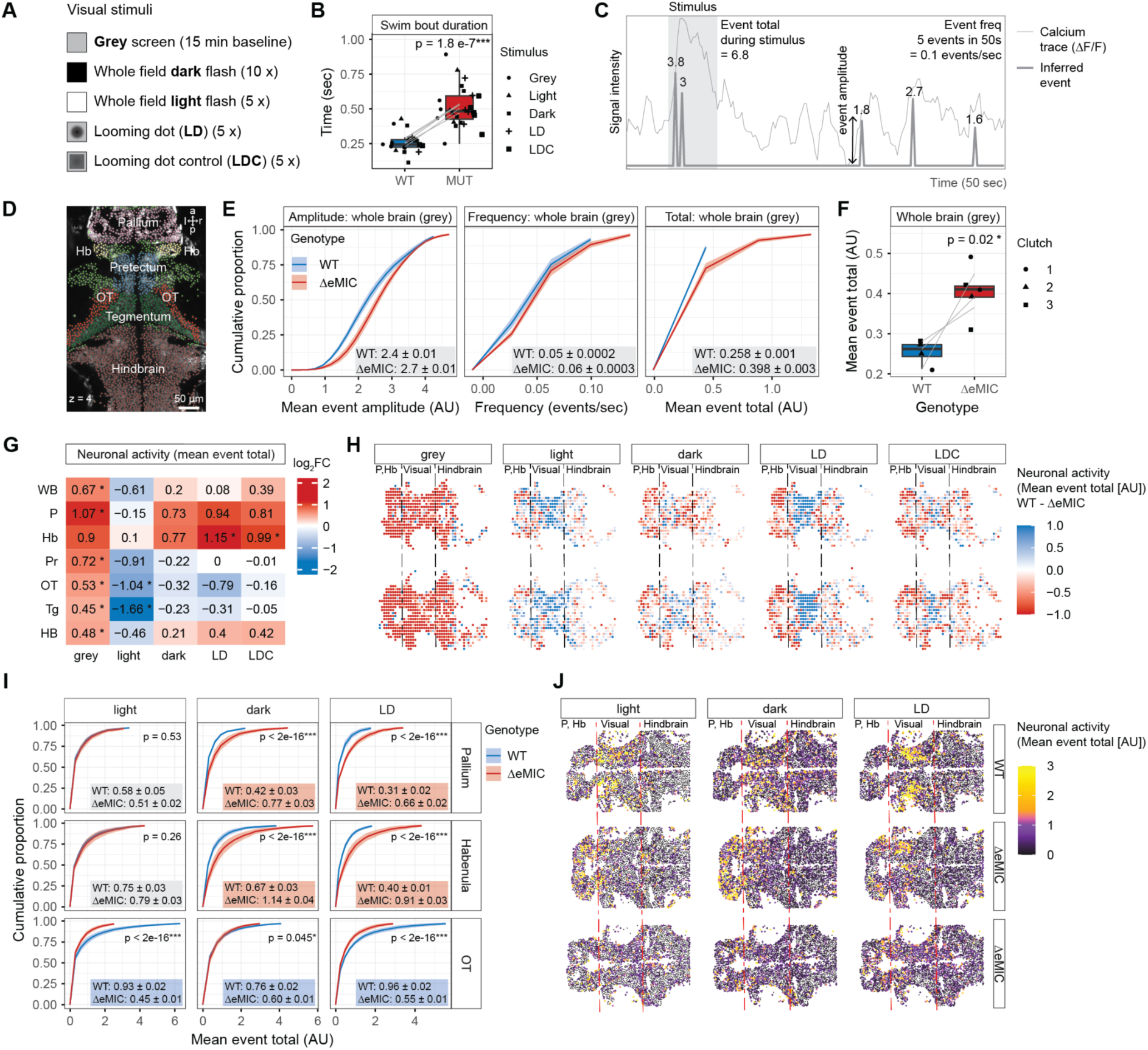
***srrm3*^ΔeMIC^ larvae show increased neuronal activity at baseline and a distinct response to visual stimuli. A:** For each z-plane, following 15 min of baseline recording, visual stimuli were presented (5 - 10 times) for ∼ 3 or 5 sec every 30 sec in a pseudo-random manner. **B:** Swim bout duration (sec) across stimuli. Each dot represents the median for one larva, shaped by stimulus. Lines connect means for each stimulus condition. Statistics by GLM controlling for stimulus and larva. n = 4 WT and 5 *srrm3*^ΔeMIC^ larvae. **C:** Diagram of a typical calcium trace (ΔF/F, thin line) and inferred spike events (thick line) with numbers indicating event amplitudes. Event total is calculated by averaging the sum of event amplitudes across same stimulus windows (grey background) and normalizing to stimulus duration (see Methods). **D:** Field of view (FOV) used for clutch 2 and 3 showing one z-plane with regions of interest (ROIs) coloured by Zebrafish Brain Browser (ZBB) anatomical region (white labels). a, anterior; p, posterior; r, rightwards; l, leftwards. **E:** Empirical cumulative distribution functions (ECDFs) of mean event amplitude (left), event frequency (centre) and mean event total (right) for the whole brain at baseline. Lines indicate means ± SEM across the same genotype and are cut at y = 0.995 for visualization. Boxed values show mean ± SEM across neurons of the same genotype. Statistics by Wilcoxon-test, comparing WT (n = 36,951 neurons) and *srrm3*^ΔeMIC^ (n = 38,940 neurons) from 4 to 5 larvae per geonotype of 3 clutches. **F:** Mean event total (AU) across the whole brain at baseline. Dots represent larva means. Lines connect means across clutches. Statistics by Wilcoxon-test comparing 4 WT and 5 *srrm3*^ΔeMIC^ larvae from 3 clutches. **G:** Heatmap of event total log2 fold-changes (log2FC) with blue indicating higher activity in WT and red in *srrm3*^ΔeMIC^ larvae. Statistics by Wilcoxon-test, comparing 4 to 5 WT and *srrm3*^ΔeMIC^ larvae from 3 clutches. Significance stars represent: *p < 0.05, **p < 0.01, *** p < 0.001. **H:** ZBB-registered anatomies of two z-planes (rows) coloured by difference in event total (AU) per spatial bin (10 µm) (see Methods). Blue indicates higher activity in the WT and red in the *srrm3*^ΔeMIC^ larvae. Values > 1 were set to 1 and all values < -1 to -1 for visualisation. **I:** ECDF of mean event totals across 3 stimuli (light, dark, LD) and 3 brain regions (Hb, P, OT). Lines indicate means ± SEM across the same genotype and are cut at y = 0.995 for visualization. Boxed values show mean ± SEM across neurons of the same genotype, with colours indicating the direction of activity change (WT higher = blue, *srrm3*^ΔeMIC^ higher = red, n.s. = grey). Statistics by Wilcoxon-test, comparing 1,407-9,451 WT - and 2,304 - 9,662 *srrm3*^ΔeMIC^ neurons per stimulus and region from 4 to 5 larvae. **J:** Anatomy plots for single z-plane of WT and *srrm3*^ΔeMIC^ larvae from one clutch with neurons (dots) coloured by mean event total (AU). Values > 3 were set to 3 for visualisation. **H,J:** Dashed lines provide anatomical guidelines dividing into left: P, Hb; centre: visually responsive (Pr, Tg, OT); left: hindbrain. **D,G,H-J:** WB, whole brain; P, pallium; Hb, habenula; Pr, Pretectum; OT, optic tectum; Tg, Tegmentum; HB, hindbrain.

For neuronal activity imaging, we generated a *srrm3*^ΔeMIC^ line with a *nacre* mutant background (without skin pigmentation)^60^ expressing a calcium indicator under a pan-neuronal promoter [*Tg(elavl3:H2B-GCaMP6s)^jf5^*]^61^. Neuronal activity was estimated from deconvolved fluorescent calcium data^62^. These deconvolved events represent calcium influx into neurons triggered by action potentials, with event amplitudes reflecting the extent of calcium entry. Here, we measured event frequency, amplitude and event total (summed event amplitudes over the stimulus window) (Fig. 3C) from neurons of the anterior dorsal brain of sibling larvae (Fig. 3D, Fig. S3B). All three parameters were significantly increased in *srrm3*^ΔeMIC^ neurons at baseline (Fig. 3E), with event frequency remaining above WT levels even when excluding periods of movement (Fig. S3C). Measuring at the level of individual larvae rather than neurons gave similar results (Fig. 3F,G; Fig. S3D,E).

Event amplitudes in the *srrm3*^ΔeMIC^ larvae remained high independently of the type of stimulus (Fig. S3F). Given similar levels of *GCaMP6s* expression levels between genotypes (Fig. S3G), this may reflect more rapid bursts of action potentials in the mutant neurons, resulting in a stronger summation of calcium transients and thus larger event amplitudes^63–65^. In contrast to the heightened neuronal baseline activity across brain regions (Fig. 3E-H), we observed generally reduced visually-evoked activity in visually-responsive brain regions (Fig. 3G-J; Fig. S3H)^66,67^, consistent with the mutants’ visual impairment^38^. This effect was most prominent during the whole field light flash stimulus, where *srrm3*^ΔeMIC^ neuron activity in the optic tectum was ∼50% below WT levels (Fig. 3I,J; p < 2.2e-16, Wilcoxon-test). Remarkably, despite this reduced response in the optic tectum, *srrm3*^ΔeMIC^ forebrain and hindbrain neuron activity remained above WT levels during stress-inducing stimuli (dark, LD) (Fig. 3G-J, Fig. S3I). Thus, in line with the behavioural data, we found that *srrm3*^ΔeMIC^ brain activity was above WT levels at baseline, and, with the exception of visually-responsive brain regions, remained increased during visual stimuli, reflecting a hyperaroused state also at the level of neuronal activity.

### Conserved programs of neural microexons are mis-spliced in *srrm3*^ΔeMIC^ neurons

We next investigated the cellular and molecular changes underlying the behavioural and neuronal hyperactivity phenotypes. To this end, we performed bulk RNA-sequencing (RNAs- seq) on fluorescence-activated cell sorting (FACS)-sorted GFP⁺ neurons from 5 dpf *Tg(elavl3:GFP)* transgenic larvae with *srrm3*^ΔeMIC^ or WT backgrounds, generating high-depth 125-nt paired end reads suitable for both alternative splicing and differential gene expression analysis (Fig. 4A). We found a marked downregulation of vision-related genes (including *opn1sw1, pde6ha, gnat2, grk7a*) in the mutants, consistent with their known photoreceptor degeneration^38^; however, these alterations may potentially mask gene expression changes driven by other neurons (Fig. S4A). To address this, we enucleated zebrafish larvae (6 dpf) prior to FACS-sorting of GFP^+^ neurons to generate both single-cell RNA-seq data and a second, shallower bulk RNA-seq dataset of 50-nt single-end reads, suitable for gene expression analysis (Fig. 4A). The latter included both dimethyl sulfoxide (DMSO)-treated larvae, used to compare gene expression profiles between *srrm3*^ΔeMIC^ and WT larvae in control conditions (Fig. 5, Fig. 6), and larvae of both genotypes exposed to pharmacological compounds (Fig. 7).

**Figure 4:**
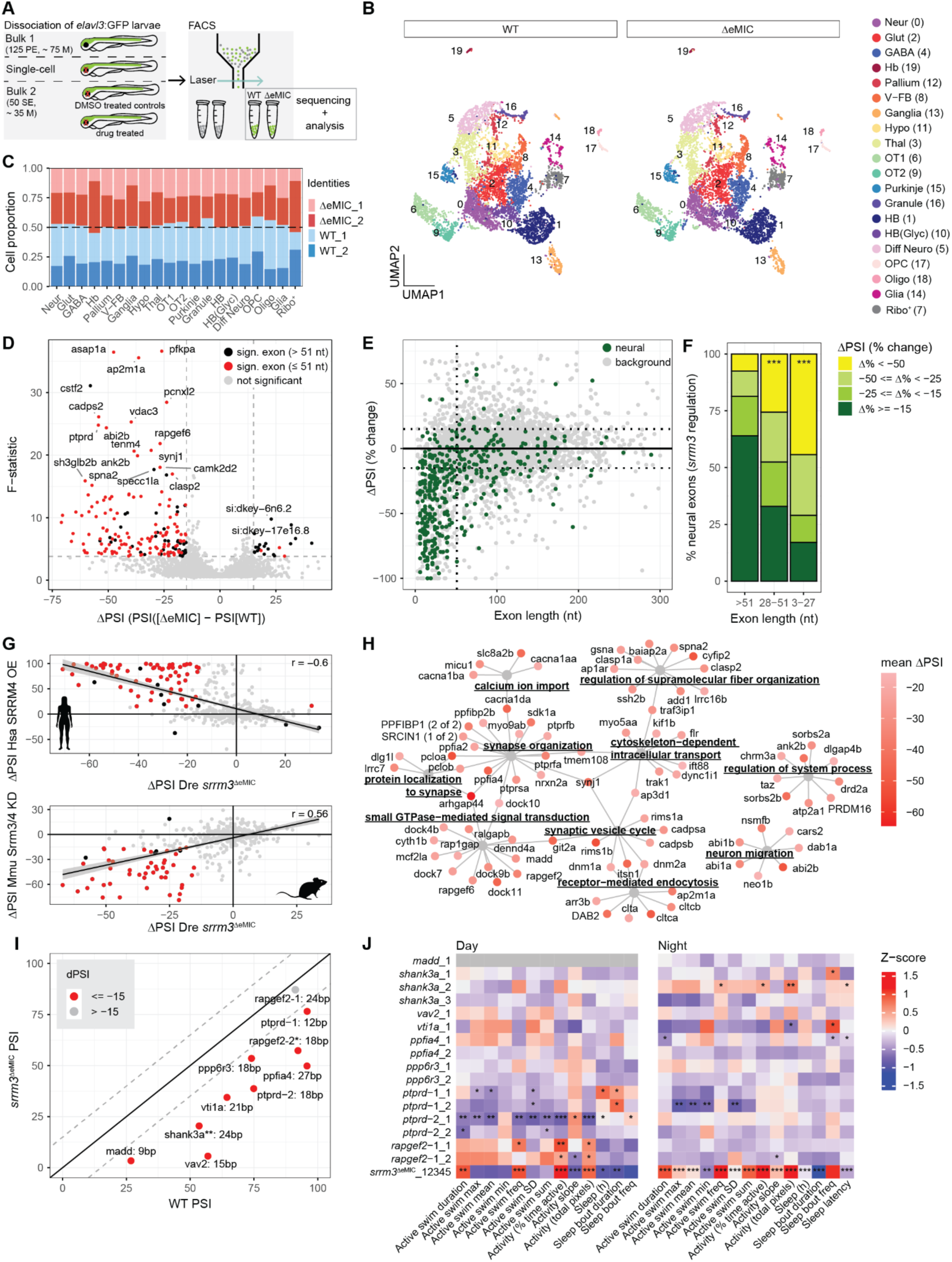
***srrm3*^ΔeMIC^ larvae show reduced microexon inclusion and no differences in cell type composition. A:** Schematic representation of the experimental design used to obtain neurons for bulk and single-cell RNA-seq (see Methods). Deep sequencing for bulk 1 (125 paired-end [PE], ∼ 75 million [M] reads) allowed for splicing analysis, while bulk 2 was sequenced more shallow (50 single-end [SE], ∼ 35 M reads) suitable for gene expression profiling. **B:** UMAP visualization of 20 single-cell clusters (numbers in order of cluster size with zero being the largest) based on PC 1-16 and nearest neighbours k = 30, split by genotype. **C:** Proportions of each genotype (WT, *srrm3*^ΔeMIC^) - clutch (_1, _2) combination contributing to the cluster compositions. Statistics with the R package *propeller*^68^**. B,C:** Cluster names: Neur, neuronal; Glut, glutamatergic; GABA, GABAergic; Hb, habenula; Pallium, pallium; V-FB, ventral forebrain; Ganglia, cranial ganglion, Hypo, hypothalamus; Thal, thalamus; OT1, optic tectum 1; OT2, optic tectum 2; Purkinje, Purkinje cells (cerebellum); Granule, cerebellar granule cells; HB, hindbrain; HB(Glyc), hindbrain (glycinergic); Diff Neuro, differentiating neurons; OPC, oligodendrocyte precursor cells; Oligo, oligodendrocytes; Glia, glia (mature, microglia, radial glia), Ribo^+^, ribosomal gene expression enriched. **D:** Volcano plot of ΔPSI (PSI[*srrm3*^ΔeMIC^] - PSI[WT]) vs F-statistic highlighting differentially spliced (F-statistic > 3.8 & |ΔPSI| > 15, shown by dashed lines) microexons (≤ 51 nt) in red and other differentially spliced exons (> 51 nt) in black. Exons not differentially spliced are in grey. Each dot represents one exon. **E:** Exon length (nt) vs change in ΔPSI (%). Neuron-specific exons^32^ shown as green - and others as grey dots. Dashed lines indicate ΔPSI (%) = -15 and 15, and exon length = 51 nt. **F:** Distribution of ΔPSI (% change) for *srrm3*-regulated neuron-specific exons grouped by length (nt). Statistics by two-sided Fisher’s exact test for exons highly skipped (Δ% < -50) vs. others between exons > 51 nts and 3 - 27 nts (odds ratio = 9.7, p.adj = 1.6e-15) and > 51 nts and 28 - 51 nt (odds ratio = 4.2, p.adj = 5e-4). P-values were Bonferroni-adjusted for multiple testing. **G:** Top: ΔPSI of conserved exons between zebrafish and human for *srrm3*^ΔeMIC^ zebrafish (PSI[*srrm3*^ΔeMIC^] - PSI[WT]) vs. SRRM4 overexpression in human 293T cells (PSI[SRRM4] - PSI[GFP control]) (r = -0.6, p < 2.2e-16, Pearson correlation)^21^. Bottom: ΔPSI of conserved exons between zebrafish and mouse for *srrm3*^ΔeMIC^ zebrafish (PSI[*srrm3*^ΔeMIC^] - PSI[WT]) vs. *Srrm3/4* knock-down mice (PSI[*Srrm3/4* KD] - PSI[WT]) (r = +0.6, p < 2.2e-16, Pearson correlation)^19^. Both: Each dot represents one exon with microexons (≤ 51 nt) or longer exons (> 51 nt) of |ΔPSI| ≥ 15 in both groups in red and black respectively. Exons with |ΔPSI| < 15 nt are shown in grey. Diagonal lines represent linear regression fit with SE. **H:** A subset of the 15 most significantly enriched Gene Ontology (GO) biological processes terms (p.adj < 0.05; terms with ≥ 5 overlapping genes excluded for clarity), for genes (dots) harbouring *srrm3*-regulated exons (ΔPSI ≤ 15) coloured by mean ΔPSI. **I:** WT PSI (x-axis) vs *srrm3*^ΔeMIC^ PSI (y-axis) of microexons for which zebrafish microexon mutant sleep/wake parameters were analysed (Fig. 4J). Dashed lines indicate ΔPSI = 15 and -15. Solid line indicates ΔPSI = 0. *No behavioural data. **Low coverage. **J:** Behavioural parameter^124,178^ heatmap for microexon mutant lines at day (5 and 6 dpf) and night (4 - 5 dpf and 6 - 7 dpf) and the *srrm3*^ΔeMIC^ (bottom). Z-scores obtained by averaging across larvae standardized to the respective WT. Grey fields indicate missing values. Number after the gene name (1 to 5) indicates biological replicate or averaged replicates in the case of 12345 for which statistics were based on all 5 clutches. Statistics by likelihood-ratio test on LMM. Detailed statistics and sample sizes are summarized in Table S1. **F,J:** Significance stars represent: *p < 0.05, **p < 0.01, *** p < 0.001.

The single-cell RNA-seq data analysis identified 20 cell clusters corresponding to distinct cell populations across anatomical regions, neural differentiation stages and major cell types (Fig. 4B; Fig. S4B,C). *srrm3* was expressed across WT neuronal clusters (Fig. S4D), in line with brain-wide *srrm3* expression as measured by hybridization chain reaction (HCR) (Fig. S4E). Comparing the proportion of *srrm3*^ΔeMIC^ and WT cells in each cluster showed a similar contribution from both genotypes (Fig. 4C; p.adj > 0.05 across clusters, t-test using *propeller*^68^), indicating no shift in cell type composition or loss of any detected population. These results are in agreement with a lack of anatomical abnormalities of the 6 dpf *srrm3*^ΔeMIC^ brain (except for reduced eye size) (Fig. S4F), pointing towards molecular alterations at the level of neuronal physiology and signalling cascades.

Next, we focussed on the detection of mis-spliced microexons in the deep bulk RNA-seq data set. Among alternative exons with sufficient read coverage (3,752; see Methods), 5% (199, in 183 genes) were significantly differentially spliced (absolute Δ percentage spliced in (|ΔPSI|) ≥ 15, F-statistic > 3.8) in *srrm3*^ΔeMIC^ neurons. As expected, the majority of these exons (89%) were skipped in the mutants, with 75% of them classified as microexons (≤51 nts) (Fig. 4D). Notably, neuron-specific microexons were also more strongly affected, as 44% of 3-27 nt exons showed > 50% reduced inclusion compared to 8% for long exons, a 5.5-fold increase (Fig. 4E,F; p.adj = 1.6e-15, two-sided Fisher’s exact test). Mis-spliced exons showed a significant overlap with orthologous exons misregulated in *Srrm3/4*-knockdown mouse cell lines^19^ (Fig. 4G; r = 0.6, p < 2.2e-16, Pearson correlation) and those affected by human *SRRM4* overexpression^21^ (Fig. 4G; r = -0.6, p < 2.2e-16, Pearson correlation), in line with the high evolutionary conservation of the *srrm3/4* microexon program across vertebrates^16,21^.

*srrm3*-dependent exons were enriched in genes involved in a range of biological processes, including “small GTPase-mediated signal transduction”, “receptor-mediated endocytosis” and “synapse organization” (Fig. 4H), in line with previous studies^16,32^. To understand the contribution of individual microexons within some of these functional categories to the *srrm3*^ΔeMIC^ hyperarousal phenotype, we performed sleep/wake analyses of zebrafish lines carrying deletions of conserved neural microexons. These include previously generated microexon mutants of *shank3*, *vti1a*, *vav2* and *madd*^32^, as well as re-analysis of published data from *ppfia4, ppp6r3, rapgef2* and *ptprd* microexon mutants^69^ (Fig. 4I,J). Interestingly, while the majority of microexon mutants showed mild to no sleep/wake phenotypes, deletion of a microexon in *rapgef2* recapitulated multiple aspects of the *srrm3*^ΔeMIC^ daytime hyperactivity, but not the behavioural changes at night. In contrast, *ptprd* microexon mutants slept more and were less active, thus showing phenotypes opposite to *srrm3*^ΔeMIC^ larvae (Fig. 4J). Taken together, while individual microexon deletions can cause sleep/wake phenotypes, it is likely that the phenotypes observed in *srrm3*^ΔeMIC^ larvae result from the combined misregulation of dozens of neural microexons.

### Upregulation of microexon-containing genes suggests a compensatory response

Differential expression analysis using the shallow bulk RNA-seq dataset from enucleated *srrm3*^ΔeMIC^ and WT larvae identified 363 differentially expressed genes (DEGs) involved in diverse neural functions (Fig. 5A; p.adj < 0.05). Downregulated genes were enriched for synaptic vesicle and endocytosis-related Gene Ontology (GO) terms, whereas upregulated ones impacted processes related to microtubules and cytoskeleton. Both DEG subsets included genes encoding calcium channels (Fig. 5B,C). Notably, multiple DEGs also contained conserved microexons with previously characterized functions (Fig. 5A,B), such as *cpeb4* (DreEX0023919: ΔPSI = -27) in neurotransmission^70^ or *micu1* (DreEX0046514: ΔPSI = -42; DreEX6103867: ΔPSI = -17) in the regulation of mitochondrial calcium levels^71,72^. This suggests that gene expression changes could be an attempt to compensate for the loss of protein function upon microexon skipping. In line with this idea, pharmacologically inhibiting the *micu1*-gated mitochondrial calcium uniporter resulted in reduced survival of *srrm3*^ΔeMIC^ larvae relative to WT siblings (Fig. S5A). To assess this potentially compensatory pattern more globally, we performed a gene set enrichment analysis (GSEA) using genes with *srrm3*- dependent microexons (ΔPSI < -15, length ≤ 51 nts) as a query set and observed a significantly strong upregulation of these genes with respect to the genomic background (Fig. S5B; normalized enrichment value [NES] = 2.6, p.adj = 1e-10). Indeed, out of 184 upregulated genes, 27% harboured *srrm3*-dependent microexons, as opposed to 1% in the background (Fig. 5D; odds ratio = 36.4, p < 2.2e-16, two-sided Fisher’s exact test). We obtained similar results in the single-cell RNA-seq data, where *srrm3*-dependent microexon-harbouring genes were significantly enriched among upregulated genes in 14 out of 19 clusters tested, including the glutamatergic (excitatory) and GABAergic (inhibitory) neuronal clusters (Fig. 5E; Fig. S5C). Given that the vast majority of *srrm3*-dependent exons - and all of the differentially included microexons in upregulated genes - are predicted to keep the protein reading frame (Fig. S5D), it is tempting to speculate that upregulation of microexon-harbouring genes occurs as a general attempt to compensate for the loss of protein function upon microexon skipping in *srrm3*^ΔeMIC^ zebrafish neurons. In line with this hypothesis, upregulation often affected the paralog of the mis-spliced gene as well, particularly if this paralog pair had a high sequence similarity (p = 0.003, Permutation test) (Fig. S5E). Altogether, *srrm3*^ΔeMIC^ larvae exhibit differential expression predominantly in neural genes important for key neurodevelopmental processes and a striking upregulation of microexon-containing genes, possibly reflecting a general attempt to compensate for a reduction or loss of protein function upon microexon skipping.

**Figure 5.**
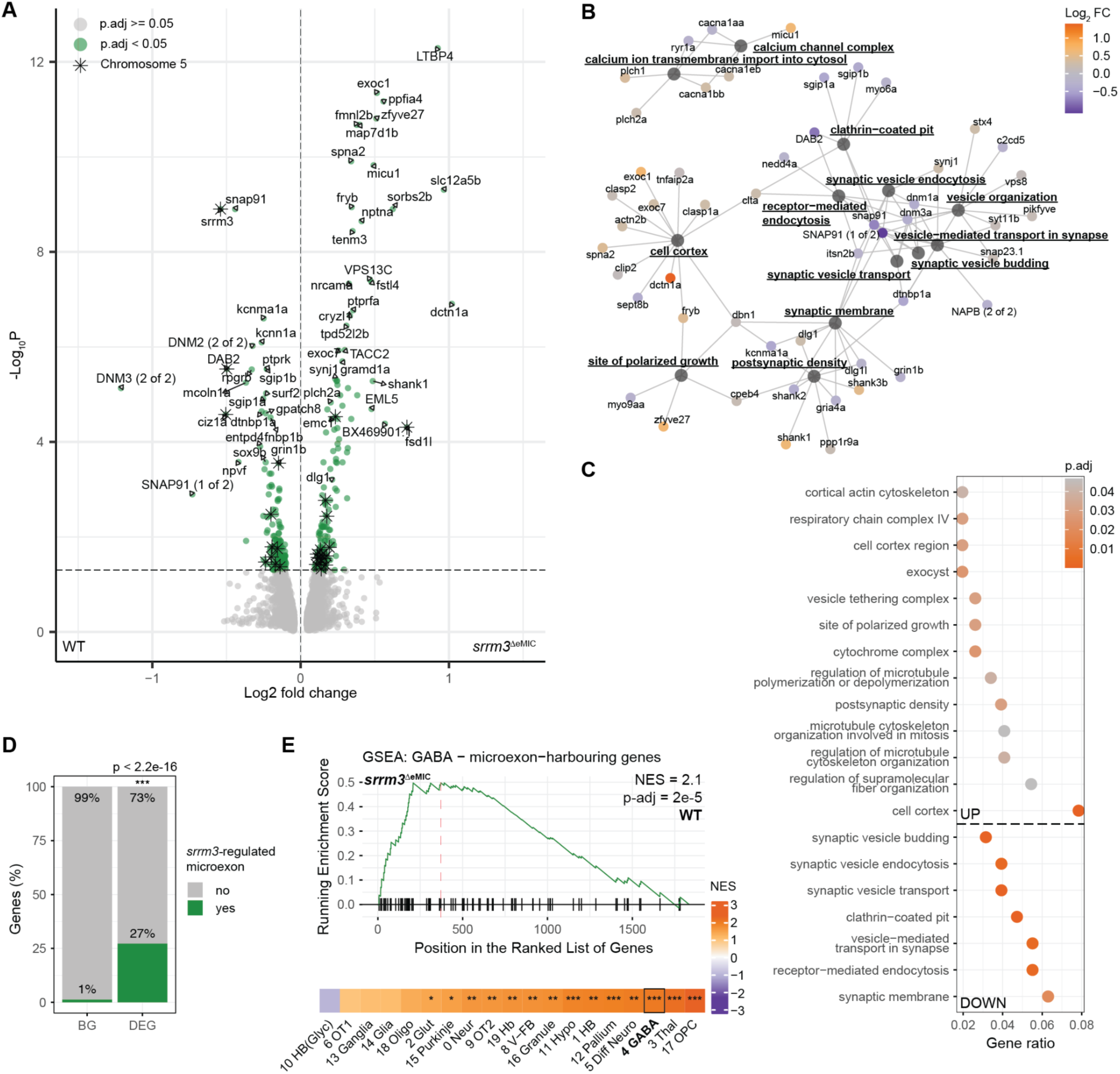
Genes upregulated in the *srrm3*^ΔeMIC^ disproportionally harbour mis-spliced microexons. A: Differential gene expression between *srrm3*^ΔeMIC^ and WT neurons of 6 dpf enucleated larvae. DEGs are shown in green (p.adj < 0.05) with those upregulated in the *srrm3*^ΔeMIC^ on the right and downregulated on the left. Stars indicate DEGs located on the same chromosome as *srrm3* (chromosome 5). N = 8 biological replicates per genotype from 10 to 24 larvae each. **B:** Enriched GO terms (p.adj < 0.05) for DEGs between WT and *srrm3*^ΔeMIC^ coloured by log2FC (orange = upregulated, purple = downregulated). **C:** Enrichment of unique GO terms (p.adj < 0.05; min set size = 10; max set size = 200) of up - (top) and down regulated (bottom) genes sorted by gene ratio and colored by p.adj. **D:** Percentage of *srrm3*-regulated microexons (ΔPSI < -15 irrespective of their F-statistic, length ≤ 51 nts) in expressed genes (background [BG], 231/18,341 genes) vs significantly upregulated genes (total = 50/184). Odds ratio = 36.4, p < 2.2e-16. Statistics by two-sided Fisher’s exact test. **E:** GSEAs for microexon-harbouring genes (length ≤ 51 nt, ΔPSI < -15) on single-cell RNA-seq data log2FC between WT and *srrm3*^ΔeMIC^ for each cluster (bottom) with an enrichment trace shown for the GABAergic population (top). P-values FDR-corrected for multiple testing across clusters. Significance stars represent: *p < 0.05, **p < 0.01, *** p < 0.001. N = 2 biological replicates per genotype with 13 to 21 larvae per replicate. Cluster names as in Fig. 4.

### Behavioural pharmacology reveals cAMP signalling as a central driver of *srrm3*^ΔeMIC^ daytime hyperactivity

Behavioural hyperactivity has been described for various zebrafish mutants, including those of genes involved in conserved intracellular signalling cascades^73,74^. Moreover, hyperactivity can also be achieved upon modulation of conserved arousal pathways^33,75^. For instance, noradrenaline acts to promote wakefulness in zebrafish^34^ and activating dopaminergic neurons can induce zebrafish locomotion^76^. To determine whether such arousal pathways are disrupted in *srrm3*^ΔeMIC^, we targeted dopaminergic and noradrenergic pathways pharmacologically and monitored the effect on daytime hyperactivity (Fig. 6A). We first targeted the dopaminergic system using the D2-like receptor agonist ropinirole. While both *srrm3*^ΔeMIC^ and WT larvae showed a significant dose-dependent reduction in activity, *srrm3*^ΔeMIC^ exhibited reduced sensitivity to the treatment (Fig. 6B; Fig. S6A). Next, we tested the response of the noradrenergic system by treating larvae with clonidine, an alpha-2 adrenergic receptor agonist. Similar to ropinirole, clonidine induced a dose-dependent reduction in activity for both genotypes, but *srrm3*^ΔeMIC^ larvae again exhibited significantly reduced sensitivity (Fig. 6B; Fig. S6A).

**Figure 6.**
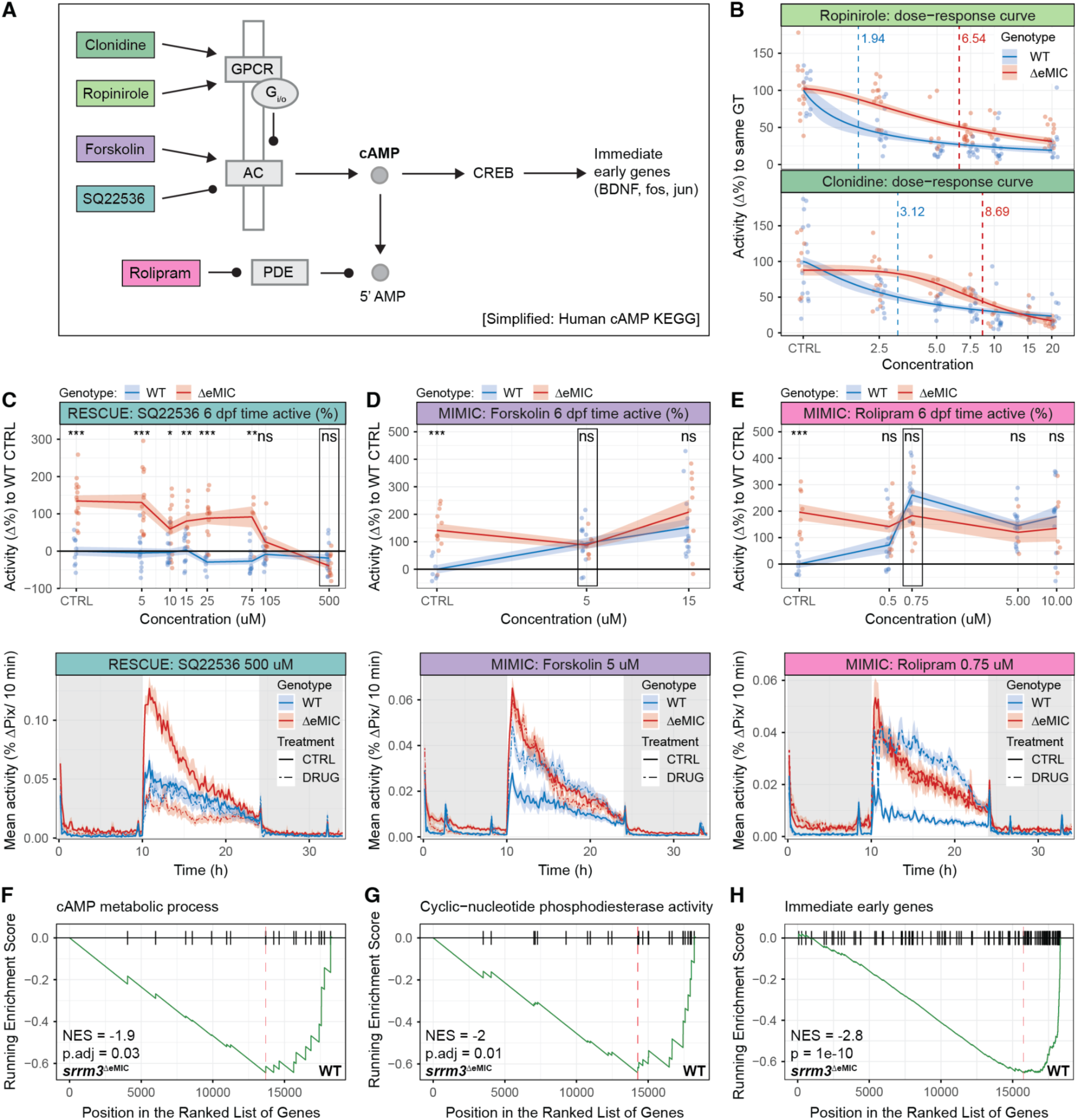
cAMP signalling is central to *srrm3*^ΔeMIC^ daytime hyperactivity. Colours across the figure correspond to each drug’s effect on cAMP levels. Green: indirectly reducing cAMP levels (clonidine, ropinirole), turquoise: directly reducing cAMP levels (SQ22563 [*srrm3*^ΔeMIC^ rescue]), pink/purple: increasing cAMP levels (forskolin, rolipram [*srrm3*^ΔeMIC^ mimic]). **A:** Simplified human cAMP KEGG pathway^179^ highlighting targets of the drugs used for behavioural pharmacology. AC, adenylyl cyclase; PDE, phosphodiesterase; GPCR, G-protein coupled receptor. **B:** Percentage change in daytime activity (% time active) of control (CTRL) and drug-treated larvae (dots) compared to same genotype (GT) controls. Traces represent the estimate ± SE of a nonlinear dose-response model fitted using a four- parameter logistic function (Hill equation) with the lower asymptote fixed at zero. Dashed lines indicate respective estimated ED50 (effective dose 50) values. Ropinirole: WT = 1.9 ± 1.3, ΔeMIC = 6.5 ± 1.2, p = 5.3e-5. Clonidine: WT = 3.1 ± 1.3, ΔeMIC = 9.7 ± 1.2, p = 1.1e-4). Concentrations on a log-scaled x-axis. Sample sizes: Clonidine: 5 to 16 larvae per group; Ropinirole: 6 to 15 larvae per group. **C-E (top):** Percentage change in daytime activity (% time active) of CTRL and drug-treated larvae (dots) compared to WT controls (black line). Concentrations on a log-scaled x-axis. Traces represent mean ± SEM across larvae from the same genotype and treatment. Boxes indicate experiments for which activity traces are shown below. Statistics by Wilcoxon-test. Significance stars represent: *p < 0.05, **p < 0.01, *** p < 0.001. Sample sizes: SQ22536 experiments: 5 to 15, forskolin experiments: 7 to 15, rolipram experiments: 4 to 16 larvae per treatment group. **C-E (bottom)**: Mean activity (%ΔPix/10 min) for either drug-treated (dashed line) or control (solid line) during 34 h (5 - 7 dpf) on a 14 h - light - 10 h dark cycle (white and grey background respectively). For visualisation purposes values for refill periods were replaced with pre refill values. Sample sizes: SQ22536 experiment: 11 to 14, forskolin experiment: 8 to 15, rolipram experiment: 8 to 13 larvae per treatment group. **F:** GSEA on log2FC between DMSO- treated WT and *srrm3*^ΔeMIC^ for “Cyclic-nucleotide phosphodiesterase activity”. **G:** GSEA on log2FC between DMSO-treated WT and *srrm3*^ΔeMIC^ for “cAMP metabolic process”. **H:** GSEA on log2FC between DMSO-treated WT and *srrm3*^ΔeMIC^ for mouse ortholog IEGs^180^.

Given that interfering with both arousal systems had similar effects, these results suggested an alteration in a common downstream pathway. Since both ropinirole and clonidine act via Gi/o-coupled G-protein-coupled receptors (GPCR) that inhibit adenylyl cyclase, thus reducing cyclic adenosine monophosphate (cAMP) synthesis^77,78^ (Fig. 6A), we hypothesized that elevated cAMP signalling might underlie the hyperactivity in *srrm3*^ΔeMIC^ larvae. Consistently, reducing cAMP synthesis with the adenylyl cyclase inhibitor SQ22536 rescued the hyperactivity in *srrm3*^ΔeMIC^, normalizing their activity to WT levels without noticeably affecting WT behaviour (Fig. 6C; Fig. S6A). In contrast, increasing cAMP synthesis with the adenylyl cyclase activator forskolin or decreasing its degradation with the phosphodiesterase inhibitor rolipram did not further increase *srrm3*^ΔeMIC^ activity but did induce hyperactivity in WT larvae, remarkably mimicking the daytime activity levels observed in *srrm3*^ΔeMIC^ (Fig. 6D,E; Fig. S6A). These results strongly suggest that elevated cAMP signalling is a central driver of the daytime hyperactivity phenotype.

To assess coordinated expression changes in cAMP signal related genes in *srrm3*^ΔeMIC^ larvae, we ran GSEA using cAMP-associated GO terms. We observed significant downregulation of phosphodiesterases (enzymes responsible for cAMP degradation) and genes associated with the GO term “cAMP metabolic process” (Fig. 6F,G; Fig. S6B). Additionally, we found significant downregulation of the zebrafish orthologs of mammalian activity-induced immediate early genes (IEGs), including *fosab*, *junba*, *bdnf* (Fig. 6H), many of which are under the regulation of the cAMP-inducible transcription factor CREB (cAMP response element-binding protein)^79–81^ (Fig. 6A). Reduced expression of the IEG *fosab* was validated by HCR-staining (Fig. S6C). While the lower expression of phosphodiesterase genes in *srrm3*^ΔeMIC^ neurons might contribute to the hypothesized elevation in cAMP levels, the observed downregulation of IEGs and “cAMP metabolic process” genes (predominantly adenylyl cyclases) likely reflects homeostatic regulation in response to sustained neuronal activity^82,83^ and prolonged activation of the cAMP pathway^84,85^.

### *srrm3*^ΔeMIC^ transcriptomic signatures are partially mimicked and reversed by cAMP pharmacological modulation

Given the ability of cAMP pathway targeting drugs to behaviourally mimic and rescue *srrm3*^ΔeMIC^ daytime hyperactivity (Fig. 6C-E), we next set out to identify common transcriptomic signatures underlying those behavioural effects. For this purpose, we compared bulk RNA-seq of FACS-sorted neurons of *srrm3*^ΔeMIC^ and WT larvae treated with drugs that increase cAMP levels (i.e., the adenylyl cyclase activator forskolin, the phosphodiesterase inhibitor rolipram, and PTZ, which indirectly raises cAMP by increasing neuronal activity) and drugs that decrease cAMP levels (i.e., the adenylyl cyclase inhibitor SQ22536, and clonidine and ropinirole, both of which activate Gi/o-coupled GPCRs thus inhibiting adenylyl cyclase) with matched DMSO-treated controls (Fig. 4A). A principal component analysis (PCA) of all samples revealed that the largest amount of variance (PC1: 17%) is explained by treatment with the adenylyl cyclase activator SQ22536 versus all other conditions (Fig. S7A). This is in line with the large number of DEGs identified upon SQ22536 treatment for both genotypes (p < 0.05; ΔeMIC: n = 3,619, WT: n = 3,829), and contrasts with the few DEGs identified for all other treatments (Fig. 7A; Fig. S7B; mean = 17 genes, range = 0 - 109 genes). SQ22536- induced DEGs were enriched for terms related to neural development (Fig. S7C), as previously reported in zebrafish^86^. Components of the adenylate cyclase-modulating G-protein coupled receptor pathway were also enriched among upregulated genes upon SQ225536 treatment of either WT or *srrm3*^ΔeMIC^ (Fig. S7C), highlighting a transcriptional effect at the level of cAMP synthesis.

**Figure 7:**
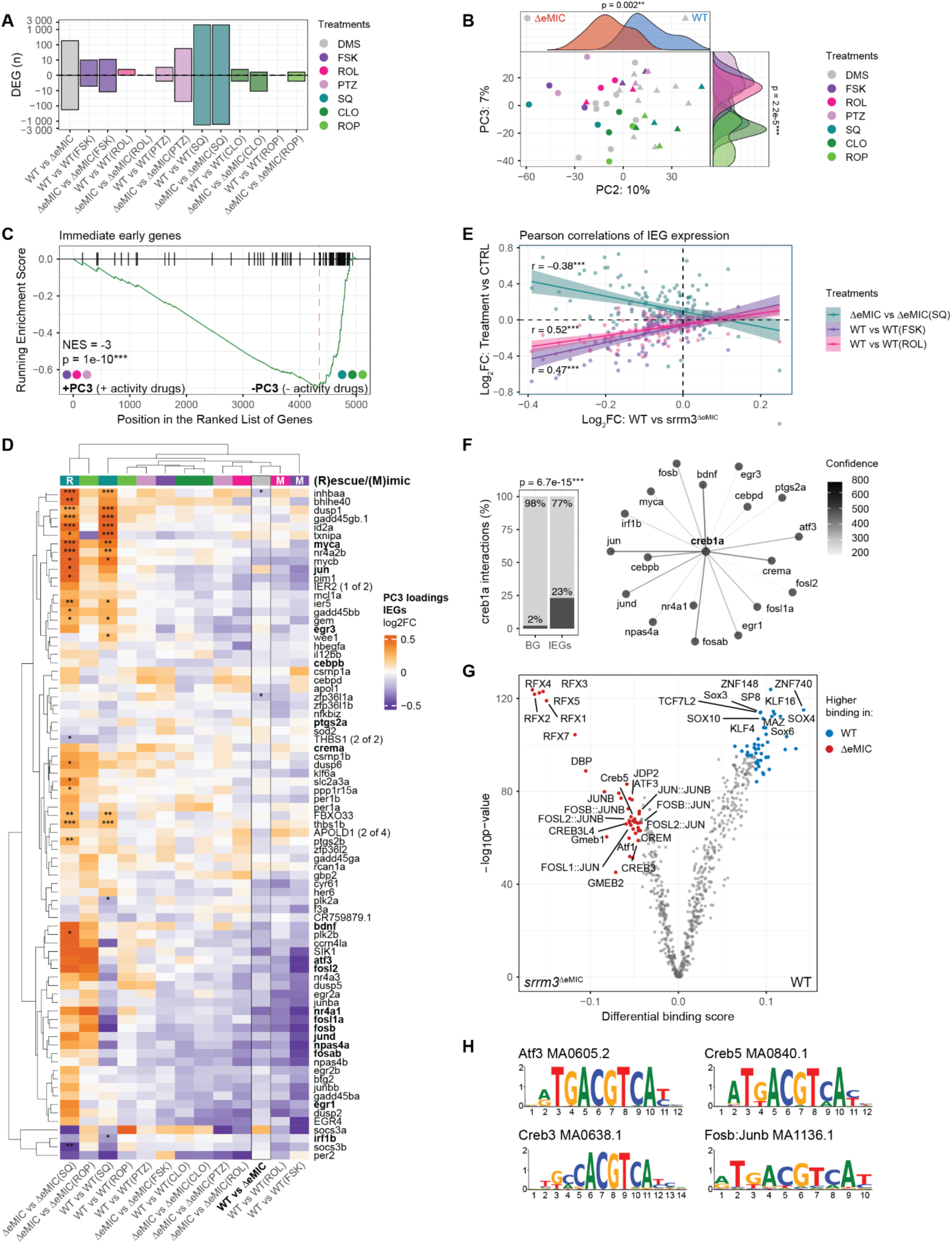
cAMP pathway modulators can partially mimic and reverse *srrm3*^ΔeMIC^-induced gene expression changes. A: Number of DEGs for each condition on a y-axis scaled using a pseudo- logarithmic transformation with a base of 10. N = 8 biological replicates for WT vs. ΔeMIC and 2 per treatment for all other comparisons but ΔeMIC vs. ΔeMIC + ROL where N = 1. Each replicate is ∼ 13 larvae per genotype and clutch. **B:** PCA on the top 5000 most variable gene counts after variance stabilisation^152^ and batch correction with “Empirical Bayes-moderated adjustment for unwanted covariates”^153^. Density ridge plots show a separation of DMSO - treated WT (blue) and *srrm3*^ΔeMIC^ (red) samples on PC2 (W = 6, p = 0.002, Wilcoxon-test) and a separation of activity-inducing (PTZ, ROL, FSK) vs. activity-reducing (ROP, SQ, CLO) samples on PC3 (W = 6, p = 2.2e-5, Wilcoxon-test). Each dot represents one sample shaped by genotype and coloured by treatment. **C:** GSEA using “Mouse ortholog IEGs”^180^ as a test set and the loadings of PC3 as a ranked background. **D:** Heatmap of IEG^180^ log2FC with colours indicating conditions (M = mimic, R = rescue [based on behavioural observations Fig. 6]). Maximum log2FC was set to 0.5 and minimum log2FC to -0.5 for visualisation. Significance stars represent: *p < 0.05, **p < 0.01, *** p < 0.001. *creb1a* interactors from panel F are highlighted in bold. **E:** Log2FC for all expressed IEGs (dots) of the WT vs. *srrm3*^ΔeMIC^ comparison (x-axis) correlate with mimic groups [WT vs. WT(FSK): r = 0.47, p.adj = 1.8e-7; WT vs. WT(ROL): r = 0.52, p.adj = 3.8e- 9] and anti-correlate with the rescue group [ΔeMIC vs. ΔeMIC(SQ): r = -0.38, p.adj = 6.9e-5] (y-axis). Linear regression lines are shown. Statistics: Pearson correlations and p-values adjusted for testing across treatments using Bonferroni. **F:** Based on the *STRING* database (v 12.0)^154^ IEGs are enriched for *creb1a* interactors (odds ratio = 17.2, p = 6.7e-15; two-sided Fisher’s exact test) (left), visualized in a network plot (right) where edges indicate the interaction strength (min = 200, max = 1000). **G:** Differential binding scores (blue = higher in the WT, red = higher in the *srrm3*^ΔeMIC^, grey = non- significant) for 750 non-redundant vertebrate transcription factor binding motifs (dots) from *JASPAR*^177^ using *TOBIAS*^176^ for all consensus sites of open chromatin (Table S3). N = 1 biological replicate from 10 to 12 larvae per genotype. **H:** A selection of motifs bound by activity-dependent transcription factors that show differential binding in panel G. **A,B,E,D:** Treatment abbreviations: CLO, clonidine; DMSO, dimethyl sulfoxide; FSK, forskolin; PTZ, pentylenetetrazol; ROL, rolipram; ROP, ropinirole; SQ, SQ22536.

**Summary figure 8:**
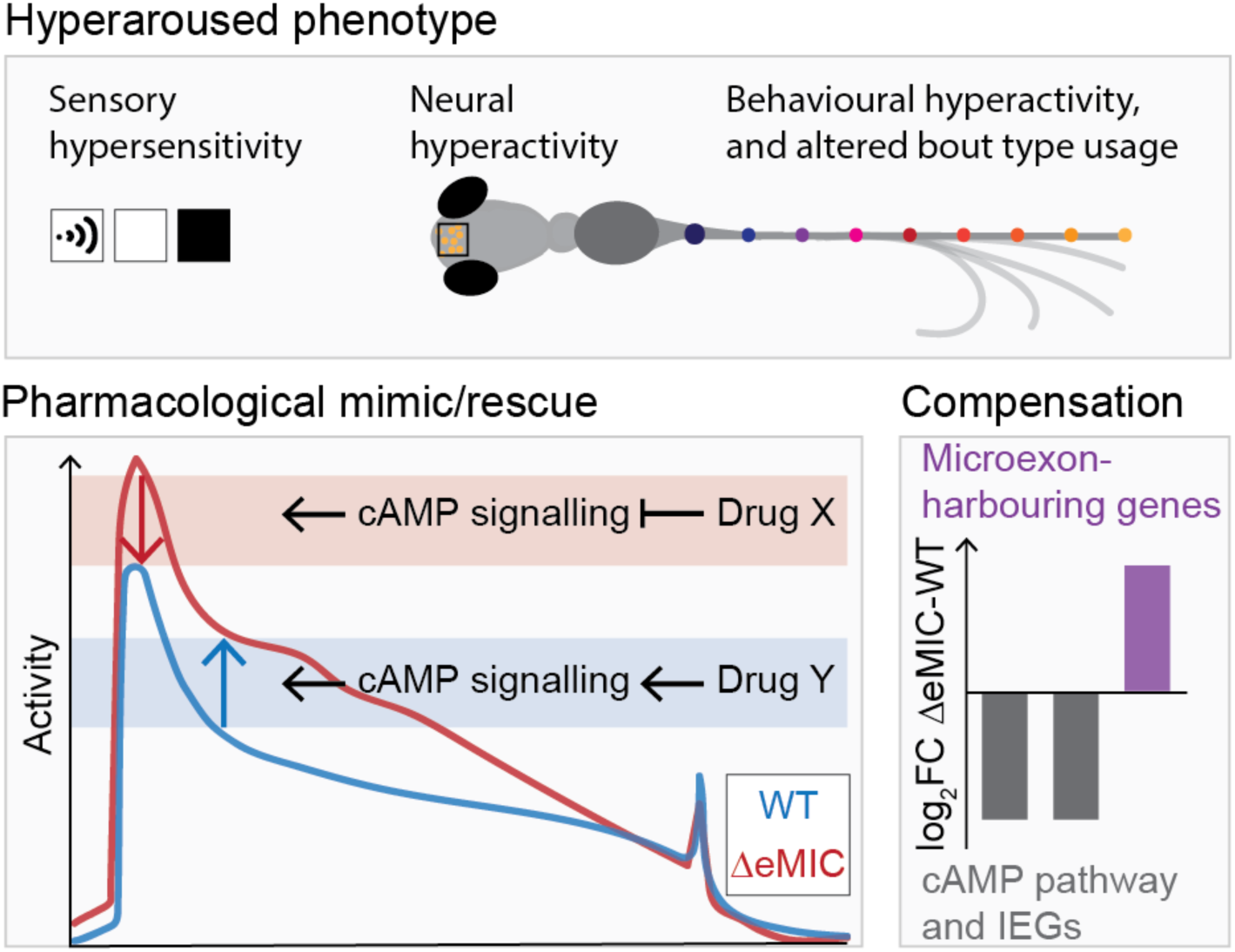
*srrm3*^ΔeMIC^ larvae exhibit a hyperaroused phenotype, reflected by increased responsiveness to sensory stimuli and heightened neural and behavioural activity at baseline. The daytime hyperactivity can be rescued upon inhibition of cAMP signalling and can be mimicked upon activation of cAMP signalling in WT larvae. Gene expression changes are suggestive of compensatory mechanisms and homeostatic regulation with microexon-harbouring genes being upregulated in the srrm3^ΔeMIC^, but cAMP pathway and IEGs, downstream of cAMP signalling, downregulated.

Beyond the substantial impact of SQ22536 (captured by PC1), we observed coherent transcriptomic patterns corresponding to treatment type. Specifically, while PC2 separated samples significantly by genotype (p = 0.002, Wilcoxon test), PC3 distinguished treatments based on their effects on daytime activity: activity-promoting drugs (forskolin, rolipram, PTZ) vs. activity-reducing drugs (clonidine, ropinirole, SQ22536) (Fig. 7B; p = 2.2e-5, Wilcoxon test). Notably, GSEA using PC3 loadings as background revealed that IEGs were significantly depleted in the treatments increasing activity (Fig. 7C). This likely reflects adaptive downregulation after prolonged stimulation (16-18h)^83,87,88^, and aligns with the downregulation of IEGs seen in *srrm3*^ΔeMIC^ larvae (Fig. 6H). The shared IEG expression profile between activity-inducing treatments (mimics; forskolin and rolipram) and *srrm3* depletion is further supported by clustering analysis (Fig. 7D) and by the significant positive correlation between IEG changes in the WT following these treatments and in *srrm3*^ΔeMIC^ larvae (Fig. 7E; Fig. S7D). Conversely, SQ22536 treatment, which mitigated *srrm3*^ΔeMIC^ daytime hyperactivity (Fig. 6C), significantly reversed many IEG alterations in the *srrm3*^ΔeMIC^ (Fig. 7E; Fig. S7D). Similar patterns were observed for genes related to “cAMP metabolic processes” (Fig. S7E), also implicating cAMP signalling in these transcriptomic responses.

Multiple IEGs encode important transcription factors, including the AP1 family members (*fos*, *jun*), whose DNA binding motifs partially overlap with CREB^89^, a key downstream effector of cAMP and major regulator of activity-induced transcription^90,91^. Moreover, an exceedingly high proportion of IEGs interact at the protein level with *creb1a* (Fig. 7F; 23% vs. 2%; p = 6.7e-15, two-sided Fisher’s exact test). Thus, we next assessed how the persistent hyperactivity of *srrm3*^ΔeMIC^ larvae affected transcription factor chromatin occupancy. For this purpose, we performed ATAC-sequencing on FACS-sorted neurons from enucleated 6 dpf *srrm3*^ΔeMIC^ and WT larvae. Notably, we found increased occupancy of AP1 motifs (Atf3, Junb, Fosb::Jun)^92^ and CREB-like motifs in *srrm3*^ΔeMIC^ compared to WT neurons (Fig. 7G,H; Fig. S7F-H) despite the reduced mRNA expression of these factors (Fig. 6H). These results are in line with findings in mice, in which AP1 factors maintained altered chromatin occupancy long after initial neuronal activation^88^. Overall, despite minimal transcriptomic changes upon treatment in all conditions except SQ22536, we were able to partially mimic and reverse the downregulation of IEG and cAMP pathway genes in *srrm3*^ΔeMIC^ by pharmacologically targeting cAMP signalling. Furthermore, an increased transcription factor occupancy at AP1 and CREB motifs suggests persistent chromatin-level alterations induced by neuronal activity.

## Discussion

Our results show that disruption of *srrm3*-dependent microexon splicing in zebrafish larvae leads to an elevated state of arousal, manifested as behavioural hyperactivity, significant sleep loss, sensory hypersensitivity, and anxiety-like behaviour. These phenotypes were accompanied by elevated neuronal activity, particularly in the forebrain, along with increased use of stress-associated high-displacement swim bouts, and occasional seizure-like episodes. Despite visual impairment^38^, *srrm3*^ΔeMIC^ larvae generally displayed a stronger behavioural response to visual stimuli than WT controls, further suggesting an intrinsic shift in arousal regulation. These results parallel findings in *Drosophila* eMIC mutants that also display sleep loss and increased seizure susceptibility^18^, pointing towards a deeply conserved role for neural microexons in shaping arousal states across species.

Sleep disturbances and sensory hypersensitivity are frequently comorbid in neurological disorders including schizophrenia and ASD^29,30,93,94^, where microexon mis-splicing is also commonly observed^16,24–26,70^. However, the contribution of microexon dysregulation to clinical manifestations of these conditions has remained largely unclear. Our findings implicate microexon mis-splicing in arousal regulation, including sleep-wake patterns but also neuronal excitability as previously proposed^24,70^. Furthermore, to date, little is known about the molecular signalling cascades linking global microexon mis-splicing, neuronal excitability and, ultimately, behaviour. Taking advantage of behavioural pharmacology, we identified the secondary messenger cAMP, common to conserved arousal systems such as the dopaminergic and noradrenergic system, as a central driver of *srrm3*^ΔeMIC^ daytime hyperactivity and possibly arousal. Specifically, we found that reducing cAMP synthesis with an adenylyl cyclase inhibitor normalized *srrm3*^ΔeMIC^ hyperactivity without affecting WT activity levels. In contrast, elevating cAMP using adenylyl cyclase activators or phosphodiesterase inhibitors induced hyperactivity in WT larvae but did not affect *srrm3*^ΔeMIC^ larvae activity. These findings align with previous research implicating cAMP signalling in the regulation of arousal and anxiety across species. Elevated cAMP levels induce hyperlocomotion in both zebrafish larvae and mice^43,95^. Furthermore, in zebrafish, forskolin treatment triggers excessive thigmotaxis^43^ and a shift from short- to long-latency startle responses^96^. In humans, dysregulated cAMP signalling has also been associated with anxiety disorders^97^. These phenotypes closely mirror those observed in *srrm3*^ΔeMIC^ larvae, including anxiety-like behaviours, hyperlocomotion, and a high usage of stress-associated long-latency escapes.

Transcriptomic analyses further revealed that cAMP pathway genes and IEGs, many of which are downstream of cAMP-CREB, including *fosab*, *bdnf* and *nr4a1*^79,80,89,98^, had reduced transcript levels in *srrm3*^ΔeMIC^ neurons. This likely reflects a homeostatic response to persistent neuronal hyperactivity and heightened cAMP pathway activation. Indeed, cAMP signalling is known to be autoregulated^99^, resulting in a downregulation of cAMP synthesising adenylyl cyclases following persistent activation^84,85^. However, in addition to adenylyl cyclases, we also observed a downregulation of transcripts of cAMP degrading phosphodiesterases, raising the possibility that low phosphodiesterase levels might contribute to cAMP pathway activation. In the case of IEGs, these genes are defined by their rapid upregulation in response to transient neural activity bursts^100,101^. Nonetheless, sustained neural activity, such as that seen in seizures^83,102^ or chronic stress^103,104^, has been reported to lead to reduced IEG expression, as observed for *srrm3*^ΔeMIC^ larvae. Interestingly, ATAC-seq revealed increased occupancy of activity-dependent transcription factor motifs, including AP1 and Creb, suggesting a shift in transcription factor engagement. This aligns with models of transcriptional priming or metaplasticity, where sustained neuronal activity leaves a molecular footprint that alters future gene responsiveness^88,105^. Supporting this, seizure-like activity in mice leads to prolonged AP1 motif occupancy and reduced subsequent induction of the IEG *c-fos*^88^, mirroring the diminished *fosab* expression we observed in *srrm3*^ΔeMIC^ larvae.

How does *srrm3*-dependent microexon misregulation lead to the observed molecular, cellular and behavioural phenotypes? Our analyses of hyperactivity and sleep-wake cycles for nine individual microexons largely showed no or mild phenotypes, in line with previous studies in zebrafish^32,69^ and the proposed role of microexons in protein specialization^106^. Thus, while a strong contribution of one or a few individual microexons cannot be excluded, our findings support a model in which the cumulative mis-splicing of multiple neural microexons underlies the observed phenotypes. Nevertheless, certain microexons can offer mechanistic insights on the observed phenotypes. For example, *rapgef2* is a key mediator of GPCR-driven cAMP signaling^107^ and contains several microexons. Mis-splicing of one of these microexons (24 nts) partly recapitulates the daytime hyperactivity seen in *srrm3*^ΔeMIC^ larvae. Elevated intracellular calcium may also contribute to increased cAMP levels, which can directly or indirectly activate adenylyl cyclases^108–110^. In line with this, using 2P calcium imaging, we observed elevated event amplitudes across brain regions and stimulus conditions for *srrm3*^ΔeMIC^ neurons, indicative of increased calcium influx. Several genes involved in calcium ion homeostasis, including *micu1,* whose pharmacological targeting leads to differential effects in *srrm3*^ΔeMIC^ and WT larvae, and *cacna1da* harbour mis-spliced microexons with previously reported effects on protein function^71,72,111,112^, suggesting that disruption of calcium signalling may substantially contribute to cAMP dysregulation and hyperarousal.

Another argument against the alternative hypothesis that misregulation of just a few microexons primarily drive hyperactive phenotypes is that *ptprd* microexon mutants in zebrafish display reduced larval activity^69^, in opposition to the hyperactivity seen in *srrm3^ΔeMIC^* larvae. In other words, misregulation of individual microexons can lead to opposite effects, as previously reported for other phenotypes (e.g. neurite growth^32^ or exocytosis^113^). Moreover, we observed upregulation of multiple microexon-containing genes, possibly as a compensatory mechanism for reduced protein function upon skipping. This may occur, for instance, when microexon skipping leads to protein isoforms with reduced protein-protein interaction strength, as seen for *ptprd* or *zfyve27*^114–116^. Thus, the transcriptional upregulation of a large fraction of microexon-harbouring genes provides indirect evidence for their functional molecular impact.

In summary, the *srrm3*^ΔeMIC^ phenotypes likely arise from the combined loss of precise microexon regulation in a wide diversity of proteins involved in processes such as vesicle transport, endocytosis, ion channel function and cytoskeletal organization. This large-scale disruption ultimately impairs key brain signalling pathways, particularly cAMP signalling. Given the recurrent mis-splicing of microexons in various neurological disorders^16,24–26,70^, also characterized by dysregulated arousal states^29,30^, future research may focus on rescuing these splicing defects. Potential strategies include indirect modulation of downstream pathways, such as targeting cAMP signalling as proposed for depression^117^, or direct correction of mis-spliced individual^118^ or groups of microexons^119^ using antisense oligonucleotides. Moreover, pharmacological modulation of microexon splicing, in part via master regulators, represents a promising therapeutic avenue^120^. Altogether, our findings uncover a novel role for neural microexons as modulators of vertebrate arousal state via cAMP signalling, offering new insight into how their dysregulation may contribute to the sensory and sleep disturbances observed in neurodevelopmental disorders.

## Materials and Methods

### Zebrafish husbandry and genotyping

Fish procedures at the PRBB were approved by the Barcelona Biomedical Research Park Institutional Animal Care and Use Ethic Committee Institutional Animal Care and Use Ethic Committee (PRBB– IACUEC). Fish procedures at UCL were approved by the UCL Animal Welfare Ethical Review Body and UK Home Office under the Animal (Scientific Procedures) Act 1986 and fish procedures at the Champalimaud Foundation by the Champalimaud Foundation Ethics Committee and the Portuguese Direcção Geral Veterinária, and were performed according to the European Directive 2010/63/EU. Zebrafish (*Danio rerio*) were grown in the PRBB facility at 28 C, or for the 2P calcium imaging experiments at the UCL fish facility at 28.5 C, on a 14 h light - 10 h dark cycle. Individual crosses were set to allow for within clutch experimental comparisons i.e. between siblings of heterozygous parents to compare WT, heterozygous and mutant zebrafish larvae. Embryos were collected, regularly screened for health and kept in E3 medium with or, for imaging experiments, without methylene blue in petri-dishes with ∼ 50 larvae per plate in a 28 C incubator on 14 h light - 10 h dark cycles (light on: 9am). Experiments were carried out between 4 dpf and 8 dpf. Sex differentiation occurs at a later age and can therefore not be reported here. Genotyping was performed as described by Lopez-Blanch *et al*.^32^ using whole larvae post-experimentally or the tip of the fin at 3 dpf allowing for full recovery^121^.

### Zebrafish lines

Experiments on WT larvae were done in the AB-line (ZFIN ID: ZDB-GENO-960809-7). Microexon mutant lines (genes and exon VastID as found in the Vast database [*VastDB*]^122^: *madd* [DreEX0044304], *shank3a* [DreEX0066086], *vav2* [DreEX0084167], *vti1a* [DreEX0084665]) and master regulator mutants were generated in-house using CRISPR/Cas9 as previously described^32,38^ and kept as heterozygous for crossing. Specifically, the *srrm3*^ΔeMIC^ was created by targeting the eMIC domain in a *Tg(elavl3:GFP)* background^123^. The main founder used for this study (referred to as *srrm3*^ΔeMIC^) is the *srrm3 crg3* line (ZFIN: ZDB-ALT-230328-5) with a 5 base pair deletion in the eMIC domain resulting in a frame shift. The second *srrm3* founder line used for the validation of the hyperactivity phenotype is the crg5 line (ZFIN: ZDB-ALT-230328-7) with a 19 base pair deletion. The *srrm3^crg3/crg3^;srrm4^crg4/crg4^* double mutant (ZFIN: ZDB-FISH-230512-7) was obtained from a heterozygous incross of *srrm3*^+/crg3^ and *srrm4*^+/crg4^ (ZFIN: ZDB-ALT-230328-6). Additionally, larvae from a *vsx1*^+/-^;*vsx2*^-/-^ incross with mutations in the homeodomain of the *vsx1* and *vsx2* transcription factors were kindly sent by the Martinez-Morales group at the Centro Andaluz de Biologia del Desarrollo (CSIC/UPO/JA), Sevilla, Spain at 48 hours post fertilization (hpf)^39^ for behavioural tracking experiments. For imaging (HCR and 2P calcium imaging) first a *nacre* line (*mitfaw2* allele)^60^ was crossed into the *srrm3^+/^*^ΔeMIC^ larvae background to avoid skin pigmentation. For calcium imaging, *Tg(elavl3:H2B- GCaMP6s)^jf5^* ^61^ were further crossed into the homozygous *nacre; srrm3^+/^*^ΔeMIC^ background. The larvae were kept in heterozygous for both *srrm3*^ΔeMIC^ and *Tg(elavl3:H2B-GCaMP6s)^jf5^* and were screened for fluorescence and genotyped at 3 dpf by fin-clipping before the experiments^121^.

### Statistics

Statistical tests are specified in the figure legends and were calculated using R (v 4.4.0). For parametric testing, we assessed normality using the Kolmogorov-Smirnov test from the R *stats* package and used log-transformations or non-parametric tests when necessary. Plots generally show p-values and/or stars as indicators of significance with the significance threshold set at alpha = 0.05. Specifically, n.s. represents p > 0.05, * represents p < 0.05, ** indicates p < 0.01, and *** signifies p < 0.001. For LMMs, statistics are specified in the text as estimate ± estimate SE; when percentage changes were given for LMMs on log-transformed dependent variables coefficients and errors were back-transformed; for Wilcoxon test the W-statistic was specified; for Fisher’s exact test the odds-ratio was given; when measures of central tendency are specified these are mean ± SE of the mean, unless otherwise specified. When applicable, p-values were adjusted for multiple testing using either Bonferroni or FDR correction as stated. When used, boxplots show the median (centre line), first and third quartiles (lower and upper box limits), and values within 1.5 x the interquartile range from the lower and upper quartiles (whiskers) plotted using the *ggplot2* (v 3.5.1) R package. Detailed statistics are provided by experiment in the methods section and also in the figure legends.

### Behavioural experiments using the DanioVision tracking system

#### DanioVision setup and tracking

Behavioural activity analysis was done using the DanioVision tracking chamber (Noldus Information Technology, Wageningen, Netherlands) and the EthoVision XT software (v15 - 17). The temperature unit was set to 28 C. Behaviour was recorded at 25 Hz. Larvae were placed singly into wells with E3. Genotypes were randomly distributed and genotyping performed post-experimentally on whole larvae^32^. 96 square-well, flat-bottom plates (Whatman™ 7701-1651 UNIPLATE™ 96-Well x 650 µl) were used for all DanioVision experiments but the thigmotaxis assay for which flat bottom circular 24-well plates (Nunc Cell-Culture treated multidishes, Thermo Scientific, Cat# 142475; 1 ml) were used. Post-experiment, larvae were poked with a P200 pipette tip to probe if they are moving. Non-responsive, physically malformed or larvae of unclear genotype were excluded. For pixel-based measures (activity = %ΔPix) the moving pixel threshold was adapted to lighting and background (18 - 20 pixels). For thigmotaxis assays larva centre-point detection was used with detection settings set to auto-detection in the EthoVision XT software.

#### Sleep/wake experiments and analysis

Larvae were plated the evening before experimental day 1 at either 4 or 5 dpf and tracked for multiple days and nights on a 14 h light - 10 h dark cycle (light on: 9am, Zeitgeber time [ZT] 0) up to 74 h. Light levels were set to 5% of the operating system (∼ 500 lux as checked with a light meter [UT382 USB, H160575608]). Evaporated E3 was topped up every morning within 1h after lights-on, unless otherwise specified. To compare the effect of *srrm3*^ΔeMIC^ on sleep/wake parameters with that of homozygous microexon mutants (genes plus VastID; common name given if applicable): *ppfia4* (DreEX0057348), *ppp6r3* (DreEX0057914), *ptprd*-1 (DreEX0002174, MeB), *ptprd*-2 (DreEX1003032, MeA6) and *rapgef2*- 1 (DreEX0060838), activity was reanalysed using the raw data of nights 4 - 5 dpf and 6 - 7 dpf and days 5 dpf and 6 dpf from Calhoun *et al.*^69^. For in-house data, parameters were calculated based on night 5 - 6 dpf and 6 - 7 dpf and for day 6 dpf using the *FramebyFrame* R package (v 1.1.0; https://github.com/francoiskroll/FramebyFrame)^124^ with prior conversion of the DanioVision to Zebrabox Viewpoint format. In brief using R, *vpSorter* was run to convert the raw data on multiple .xls files into one big .csv followed by *ggActivityTraceGrid,* to quality-check traces and day/night boundaries. The *multiBehaviourParameter* function was used to extract the parameters for downstream analysis and *calculateFingerprint* was used to obtain z-scores for the heatmaps. Statistics were run using *LMEreport* running a likelihood-ratio test on linear mixed effect models: *lmer(values ∼ Group/Genotype + (1|Clutch), data = night or day)* (Table S1). Percentage changes were calculated based on model estimates, where the y-intercept represents the estimated WT mean and the WT mean plus the estimate represents the ΔeMIC mean. The formula used was *(100*(ΔeMIC-mean/WT-mean))-100* if the WT- mean was positive and *((100*(ΔeMIC-mean/WT- mean))-100)*-1* if the WT-mean was negative^124^. For plotting of activity/sleep traces over time water refill periods were excluded and replaced with values of the equal amount of minutes before the water refill instead.

#### Photomotor Assay

Larvae were plated in the morning and left to acclimatise for 1.5 h to the DanioVision chamber before being subjected to first 1 h of dark and then 1h of light (100%, ∼ 10,000 lux), alternating periods to add up to 10 h total. Responsiveness (mean difference) was calculated by subtracting activity (%ΔPix) 1 min before the stimulus from activity (%ΔPix) 1 min after the stimulus per transition and then averaged for each larva. Statistics by maximum likelihood test on LMM using the R package *lme4* (v 1.1-36)^125^ with *lmerTest* (v 3.1.3)^126^: *lmer(mean difference ∼ Genotype + (1/Clutch)).*

#### Mechanical tapping habituation assay

During the day, following 1 h of acclimatisation to the DanioVision chamber, larvae were exposed to 10 * 90 sec inter-stimulus-interval (ISI) tapping stimuli [pre-habituation condition] followed by 100 * 5 sec ISI stimuli [habituation condition] at maximum tapping power (8). Larvae responding to ≥ 7/10 pre-habituation tapping stimuli were used for the analysis. Larvae were termed responsive if activity (%ΔPix) 1 sec after tapping was greater than 1 sec before tapping adapted from Fitzgerald *et al.*^127^. A habituation index (HI), based on Pantoja *et al.*^128^ was computed by calculating the difference in response probability for the last 40 stimuli (low ISI) to the response probability during the first 10 stimuli (high ISI) *(HI = 1- (Plast40/Pfirst10))*. Statistics by maximum likelihood test on LMM using the R package *lme4* (v 1.1- 36)^125^ with *lmerTest* (v 3.1.3)^126^: *lmer(HI ∼ Genotype + (1|Clutch), data = data)*.

#### Thigmotaxis assay

Experiments were performed in 3 technical replicates across the day (between ZT1 - ZT9) to maximise sample size (3 * 24 wells). Following 30 min of acclimatisation at 100% light (∼ 10,000 lux) of the operating system, larvae were recorded for 1 h. Measurements were in mm and sec. Larvae with a pearson correlation < 0.9 between activity (%ΔPix) and distance moved (mm) were discarded as this suggests tracking errors (n = 23/144 larvae). Each well was split into centre (radius [r] = 5.63 cm) and periphery (r = 8 cm) with an equal area of ∼ 100 cm^2^ per arena. Thigmotaxis (% TDM) was computed as: TDM in the periphery divided by TDM in the whole arena, multiplied by 100. Statistics by maximum likelihood test on LMM using the R package *lme4* (v 1.1-36)^125^ with *lmerTest* (v 3.1.3)^126^: *lmer(thigmotaxis ∼ Genotype + (1|Clutch), data = data)*.

### High resolution tail tracking and bout type analysis

#### Experimental design and set-up

Experiments were carried out in the Orger lab at the Champalimaud Centre for the Unknown in Lisbon, Portugal. Larvae from a *srrm3*^+/ΔeMIC^ incross were shipped at 24 - 48 hpf, raised in groups of ∼ 100 on large petri dishes (r = 14 cm) in a 10 h light - 10 h dark incubator at the local facility. Behavioural tracking was performed for 5 and 6 dpf larvae from four clutches: C1-C4. The tracking setup was described previously^50,52^. In brief, larvae were tracked in an acrylic arena with a rounded edge (radius of curvature: 11 mm, diameter: 10 cm, depth: 4 mm) (Fig. 2A) and illuminated from below using an infrared light. The arena was sheltered from outside lighting. The larvae’s eyes and tail (nine equally spaced segments of 300 μm) were tracked online at 700 Hz throughout the experiment via custom C# software and swim bouts were detected as in Marques *et al.*^50^. Experiments were performed blindly as larvae were genotyped post-tracking^32^. Larvae were left to acclimatise to the arena for 5 min, followed by a 25 min tracking period under constant illumination at ∼ 1,500 lux (C1-4) and alternating 5 min periods of darkness and light (∼ 1,500 lux) (C1,3,4). Alternatively, some (C1, C2) were tracked under light-on (∼ 1,500 lux) conditions for 15 min exposed to 10mM PTZ [P6500-25G, Sigma; stock = 100mM in E3].

#### Data pre-processing and swim bout categorization

Data was pre-processed in *MATLAB* as previously described^50,52^, obtaining the following output files: “bouts.mat” with information on bout category, bout kinematics and PC coordinates; “camlog.mat” with frame-by-frame information on the tail tracking angle across segments; “stimlog.mat” with information on light/dark stimuli. Relevant information was extracted in *Python* and summarised into data frames, one with bout data, analysed in R and one with tail angle data analysed and visualised using *Python*. Bout categorization was carried out by assigning each bout to 1 of 13 bout types based on k nearest neighbours (k = 50) analysis using an existing balanced data set of labelled examples of each bout type^50,52^. Each bout type was described by 73 kinematic parameters and embedded into a previously computed PC-space^50^ based on a large set of behavioural data under various stimulus conditions. This space and CoM of bout clusters were used for the PCA to visualise bout distributions and to calculate Euclidean distances to the CoM for each bout category in *MATLAB*. Euclidean distances were compared using Wilcoxon-test and p-values FDR adjusted for the number of bout categories tested (Table S2).

#### Bout type usage

To reliably compare bout type usage, larvae with low bout frequency of < 5 bouts/min during either baseline, dark or light periods were removed (n = 2 across 4 clutches). Percentage usage was calculated for baseline (25 min, N = 4 clutches) and dark (15 min total, N = 3 clutches) per larva. Statistics by maximum likelihood test on LMM on log-transformed counts using the R package *lme4* (v 1.1-36)^125^ with *lmerTest* (v 3.1.3)^126^: *lmer(log(count) ∼ Genotype + (1|Clutch), data = data)*. Percentages and percentage standard errors provided in the text were calculated from log-linear model^129^ estimates using the formulas: percentage = *(exp(estimate)-1)*100*; percentage SE = *exp(estimate) * estimate SE * 100.* P-values were Bonferroni-corrected for the number of bout types tested per condition (Table S2).

#### Dark-light escape response latencies

For escapes during light-dark transitions (C1, C3, C4: n = 3 transitions) larvae with no bouts in either of the 5 min dark or light periods (ΔeMIC n = 1, +/ΔeMIC n = 1) were excluded. The mean escape latency was computed by measuring the duration (ms) until the first escape (O-bend, SAT, LLC, SLC) within a 3 sec window after the stimulus. Statistics by maximum likelihood test on LMM on log-transformed counts using the R package *lme4* (v 1.1-36)^125^ with *lmerTest* (v 3.1.3)^126^: *lmer(log(latency) ∼ Genotype + (1|Clutch), data = data)*. Percentages and percentage standard errors provided in the text were calculated from log-linear model^129^ estimates using the formulas: percentage = *(exp(estimate)-1)*100*; percentage SE = *exp(estimate) * estimate SE * 100.*

#### Bout kinematics

Bout duration was computed by subtracting bout start times from bout end times. Displacement was calculated as *sqrt((boutDistanceX)^2^ + (boutDistanceY)^2^)*. Speed was calculated as *sqrt((boutSpeedX)^2^ + (boutSpeedY)^2^)*. *meanBoutFreqCorr* was computed by dividing the number of detected half-beats by the duration of the given bout. *boutAngle* encodes changes in the larva’s heading angle from the beginning to the end of the bout. The *boutMaxAngularSpeed* represents the maximal change in tail angle during the bout ^50^. Bouts with *boutAngle* > |360| C) were removed as they likely represent tracking errors (n = 20/600,469). Only bouts with values for all parameters tested (*boutMaxAngularSpeed, boutDuration, boutAngle, displacement, speed, meanBoutFreqCorr*) were included in the downstream analysis (excluded = 1,530/600,449). For statistical comparison of kinematic parameters by bout type, measures for all larvae with < 3 bouts for a given bout type were also excluded (BS: 12/190 larvae; HAT: 3/194 larvae; LLC: 4/194 larvae; O-bend: 18/178 larvae; SAT: 5/190 larvae; SLC: 23/182 larvae) (Table S2). For statistical comparisons absolute values were obtained. First, bout kinematic means across all bout types were compared using a maximum likelihood test on LMM using the R package *lme4* (v 1.1-36)^125^ with *lmerTest* (v 3.1.3)^126^: *lmer(mean ∼ Genotype + (1|Clutch), data = data)*. Similarly, kinematic parameters were compared within each bout type individually [*lmer(values ∼ Genotype + (1|Clutch), data = data)*] and Bonferroni corrected based on the number of bout types tested. For the latter difference (%) values were plotted using the *ComplexHeatmap* R package (v 2.20.0) calculated as *(mean ΔeMIC - mean WT/ |mean WT|)*100)*.

#### Long bout analysis

Long bouts were defined as > 500 ms. All larvae and experimental time points were considered in this analysis. To distinguish artefacts/tracking errors from real long bouts, tail angle traces were plotted using *Python* and manually labelled as either noise/unclear or long bout. The experimenter was blinded to genotype. For statistics a p-value was simulated by counting the number of times (out of 1000) that an equally sized set of bouts from the same genotype showed a value more extreme than that of the set of long bouts. To avoid zero-probability estimates, the p-value was calculated as *(r + 1)/(n + 1)*, where *r* is the number of permutations more extreme than the observed value and *n* is the total number of permutations^130,131^. For example, if the long bouts have a mean absolute bout angle of 67 degrees and out of n = 1000 randomly sampled sets the mean bout angle was found to be higher for r = 5 permutations, then the probability of observing, a mean absolute bout angle of > 67 degrees, by chance would be estimated at *(5+1)/(1000+1) = 0.006*.

### 2P Calcium imaging

#### Set up and experimental design

Experiments were conducted in the Bianco lab at UCL in London, UK. The custom-built 2P imaging setup and experimental procedure were described by Antinucci *et al.*^132^. Briefly, larvae were embedded in 3% low-melting agarose with eyes and tail free to move the afternoon prior to the imaging day at 5 or 6 dpf. For functional imaging the sample was illuminated at 920 nm with a laser power of ∼ 11.2 mW and for anatomical stacks at 800 nm with a laser power of ∼ 9.1 mW. Data was obtained for a total of 9 WT and *srrm3*^ΔeMIC^ larvae [nacre; *Tg(elavl3:H2B-GCaMP6s)^jf5^*] with N = 3 clutches (C1-3) with 1 - 2 larvae each per genotype. For C1 images were taken at a resolution of 500 x 469 pixels with pixel size of 0.7 µm. For C2 and C3 image resolution was 693 x 368 pixels with a pixel size of 0.61 μm to image more of the pallium and hindbrain (Fig. 3D; Fig. S3B). The FOV includes parts of the pallium, habenula, pretectum, optic tectum, tegmentum and anterior hindbrain (Fig. 3D; Fig. S3B). Acquisitions were made at 3.6 Hz for 3 - 4 planes per larva with 10 μm z-spacing, scanned one plane at a time. Visual stimuli were designed and presented as described in Antinucci *et al.*^132^ but the order was altered. Namely, 30 x grey screen epochs (each 108 frames) followed by a pseudo-random presentation of 4 different stimuli every 30 sec, including 5 x whole-field light (∼ 3 sec), 10 x whole field dark (∼ 3 sec), 5 x looming dot (∼ 6 sec), 5 x overall dimming as control for the looming dot (∼ 6 sec)^58,133^. Simultaneous tail tracking allowed for behavioural read-outs^132^.

#### Image registration

The 3D registration to the ZBB^134,135^ was performed as previously described^132^ using the ANTs toolbox (v 2.3.5)^136^ with a multi-step registration. Functional image volumes were first registered to a larger, high-resolution volume (anatomy stack) acquired of the same larva at 1 µM z-spacing. The anatomy stack was prior registered onto the “huC_nls_mCar1.nrrd” ZBB reference (1 x 1 x 1 xyz μm/px), thus transformation could be concatenated to bring the functional imaging volume onto the ZBB atlas.

#### Calcium imaging analysis: pre-processing

Functional imaging data was processed as described in Antinucci *et al*.^132^ using a standardized *MATLAB* pipeline. Motion-correction^137^ resulted in overall good alignments (> 75 % per plane) with the exception of one *srrm3*^ΔeMIC^ larva (clutch 1) with only 40% of well-aligned epochs, thus excluded from all but baseline (grey screen) analysis, for which good alignments were available. Extracted ROIs^138^ were overlaid with anatomical masks to assign cell identities. ROIs were included if localized within the *elavl3* brain mask (i.e. neurons) and subsequently grouped into anatomical regions: optic tectum (optic tectum - neuropil, optic tectum - *stratum periventriculare*), hindbrain (*corpus cerebelli*, *valvula cerebelli*, hindbrain), pretectum, tegmentum, habenula, pallium. ROI grouping was adjusted manually when necessary.

#### Swim bout kinematics in head-restrained larvae

Bout frequency was calculated per epoch spanning around 30 sec with stimuli presented within the first 3 - 5 sec of these epochs. Both bout frequency and median bout duration were based on the whole imaging session for each larva, including periods for which neuronal activity data was discarded due excessive motion. Statistics by GLM using the *glm* function from the stats R package (v 4.4.0): *glm(parameter ∼ Genotype + stimulus + ID).*

#### Calcium imaging analysis: event amplitude, frequency and event total

Neuronal activity was estimated from deconvolved fluorescent calcium data using *OASIS*^62^ in *MATLAB* using inferred spike trains (referred to as events) as input for downstream analysis in R. Since spike inference can vary depending on signal-to-noise ratios, *GCaMP6s* sensor brightness (influenced for example by expression level), was measured, when excited at 800 nm (fluorescence independent of neuronal activity)^139^ by calculating the mean grey value in *Fiji/ImageJ2* (v 2.14.0/1.54f) across multiple ROIs in the pallium (n = 2 planes with 2 measures each), habenula (n = 2 planes with 2 measures each) and optic tectum (n = 3 planes with one measure) as representative regions (Fig. S3G). For subsequent analysis of inferred spikes (events), ROIs with event amplitudes > 99 were excluded from the analysis, likely representing artefacts as most events are expected to lie between 3 and 5^62^. Analysis windows for event amplitude, frequency and event total spanned a minimum of 2 repetitions per stimulus or ∼ 6 min for grey screen baseline. The <u>mean event amplitude</u> (AU) per ROI was computed by averaging amplitudes of all non-zero events during the same stimulus (light, dark, LD, LDC) or during no stimulus grey screen periods. The <u>event frequency</u> (events/sec) per ROI was obtained by counting non-zero events during the same stimulus (light, dark, LD, LDC) or during no stimulus grey screen periods, divided by the total number of frames and multiplied by 3.6 Hz (acquisition rate) to obtain events per second. The <u>mean event total</u> (AU) was computed by summing events across each stimulus window (light, dark: 12 frames; LD, LDC: 21 frames; grey: 108 frames) and dividing by the number of windows to obtain an average for each ROI. Values were then normalized by stimulus duration to that of one light/dark epoch (12 frames, ∼ 3 sec). Statistics were performed using Wilcoxon-tests at the level of individual ROIs/neurons for each region and stimulus but also at the level of averaged values per larva for each stimulus.

#### Calcium imaging analysis: visualization

ECDFs were computed with the R *stats* package, first for each larva individually and then averaged on a common grid of x-values, based on minimum and maximum parameter values for a given stimulus condition and brain region. Lines were displayed as mean ± SEM and cut at y = 0.995, to visually capture 99.5% of the values. For visualization of mean event total in the ZBB space (Fig. 3H), ROIs were binned based on coordinates falling within 10 µm along the x, y, z axis. For example, ROIs that fall within the coordinates of x = 10 - 20, y = 10 - 20 and z = 10 - 20 were assigned a new coordinate of x = 15, y = 15 and z = 15 with the median value across ROIs. For each bin the mean across neurons from the same genotype was computed and neurons with < 3 ROIs per bin and genotype were removed. Final values for the anatomy plots were obtained by subtracting *srrm3*^ΔeMIC^ means from the WT mean.

### Bulk and single-cell RNA-seq

#### Sample preparation and dissociation

Information on each sequencing experiment can be found in Table S4. The bulk RNA-seq data for the splicing analysis was collected from 5 dpf larvae that were not enucleated. For all other sequencing runs 6 dpf larvae were used, to age-match the sleep/wake behavioural analysis, and enucleation was performed under anaesthesia (∼14 drops of tricaine into petri dish [0.4 g/100 ml]) at the day of dissociation, to avoid photoreceptor degeneration masking gene expression changes driven by other neurons (Fig. S4A)^38^. The dissociation was performed in a sterile environment following Lopez-Blanch *et al*.^32^. In brief, after a wash in Neurobasal^TM^ Media (NB) medium (Thermo Fisher - 21103049) larvae were suspended in 350 µl of NB media with supplements (1% Penicillin Streptomycin, 1% N2, 1% L-Glutamine, 2% B-27 without vitamin A) plus 125 - 150 µl of 0.5% trypsin-EDTA for enzymatic dissociation, aided by three rounds of mechanical dissociation using pestles, lasting 5, 4 and 3 min. The trypsinization reaction was halted by adding 800 µl of NB + supplements (+2% fetal bovine serum [FBS]) and the mix was spun down for 4 min at 5000 revolutions per min (rpm). The supernatant was removed and the cells resuspended in 500 µl NB with supplements and 2% FBS for the bulk experiments and in 1 x distilled phosphate-buffered saline (PBS) for the single-cell experiments via careful pipetting. The cell suspension was passed through a 40 µm filter (Cultek - 88141378) and kept on ice until FACS.

#### FACS

Neurons in *srrm3*^ΔeMIC^ larvae with *Tg(elavl3:GFP)* background were FACS-sorted. Prior to FACS propidium iodide (PI) was added (1:1000). Cells were sorted for green fluorescence (GFP - 488 nm) using the BD influx machine at a sort mode of 1.5 drop pure with nozzle size 100 µm. Following Lopez-Blanch *et al.*^32^, gatings were adjusted to filter out debris, aggregates and dead cells before selecting for up to 500,000 GFP-positive cells that were collected in Eppendorf tubes with 300 µl NB plus supplements (see “Sample preparation and dissociation”). The percentage of alive cells was on average 75% per sample (Table S4). For the single-cell experiment samples were FACS-sorted into a 96-well plate (to avoid sticking to tube walls) with 3.7 µl of 0.378% bovine serum albumin (BSA) in 1 x PBS for a final concentration of 0.04% BSA to minimise cell loss and aggregation. Using a P200 (LabClinics), cells were then transferred into low binding tubes (525-0130, VWR).

#### RNA extraction prior to bulk RNA-seq

The RNA of the sorted GFP-positive neurons was extracted in one of two ways. A) Using trizol for the splicing analysis experiment: Samples were spun down for 3 min at 5000 rpm, the supernatant removed, suspended in 500 µl of Trizol (Cat# 15596018, Thermo Fisher) and frozen at -80 C for at least 1 h. RNA extraction was performed following the instructions for Phasemaker^TM^ Tubes (Invitrogen, Cat. A33248). In brief, cell lysis was enhanced by bead beating (MiniBeadBeater-16, Biospec Products) 2 rounds of 30 sec with a 5 min break at 4 C. RNA was precipitated with an additional 2 µl of glycogen as carrier, then solubilized and finally resuspended in RNase-free water. B) PicoPure Mini RNA isolation kit for differential gene expression analysis samples: (ThermoFisher, Cat:KIT0204): RNA was extracted following the PicoPure manual. First, RNA was extracted from pellets: in brief samples were centrifuged, cells resuspended in the extraction buffer, incubated for 30 min at 42 C and again centrifuged to obtain the supernatant for downstream processing. Second, the RNA was isolated: in brief, columns were pre- conditioned, samples were mixed with equal volume of 70% EtOH, and RNA was bound to the column via centrifugation. Columns were washed with a wash buffer via centrifugation. Thirdly, purification columns were treated with DNase using the RNase-Free DNase Set (Quiagen, cat# 79254) for 15 min at room temperature (RT), washed further and RNA was eluted twice using the elution buffer into a total volume of 30µl.

#### Bulk RNA-seq

The eluted RNA was sent for library preparation to the CRG Genomics facility using the SMART-Seq v4 Ultra Low Input RNA kit in combination with the NEBNext Ultra II kit with RNA of 20 - 50 ng. For splicing analysis samples were sequenced paired-end with 125 base pair read length, generating an average of ∼ 75 million reads. Drug/DMSO-treated samples were sequenced single-end with 50 base pair read length. Sequencing was done at NextSeq2000, with an average of ∼ 35 million reads per sample (Table S4).

#### Single-cell RNA-seq

Samples were sequenced at the CRG Genomics facility using the Next GEM Single Cell 3’ Reagent Kits v3.1 on a NextSeq 500 platform (28-8-0-91) with paired-end 150 bp reads. Sequencing was performed using 150 cycles, covering either 66% (batch 1) or 100% (batch 2) of a high-output flow cell. Samples were provided in 37.5 µl of cell suspension (1x PBS, 0.04% BSA) from freshly FACS-sorted cells with a concentration of 345 cells/µl and a total number of 13,000 cells for a target recovery of 9,000 cells.

#### Single-cell RNA-seq data processing

Sequencing data was processed using *cellranger* (v6.1.2, 10X Genomics). A gene transfer format (GTF) file filtered for protein-coding genes (attribute: gene_biotype=protein_coding) was generated using the *cellranger mkgtf* and a reference was generated using *cellranger mkref* for the *Danio rerio* (zebrafish) genome assembly version GRCz10.9. Reads were aligned to the *Danio rerio* GRCz10 reference genome using *cellranger count*, which also performed unique molecular identifier (UMI) counting for gene expression quantification. Introns were included in the analysis, and the expected number of recovered cells was set to 9,000.

#### Single-cell clustering and downstream analysis in R

A total of 4,024 (WT 1), 3,200 (WT 2), 2,946 (*Δ*eMIC 1) and 4,070 (*Δ*eMIC 2) cells were obtained from sequencing with 145 - 172 million reads per sample. The total 14,240 single-cell transcriptomes were further quality-filtered and used for the single-cell analysis. Cells were filtered based on average values obtained from the *isOutlier* function (median absolute deviation > 3) from the *scater* R package (v 1.32.0) across replicates: > 300 expressed genes, < 5% mitochondrial content, > 400 and < 30,000 UMI count. A total of 3,836 (WT 1), 2,808 (*Δ*eMIC 1), 2,785 (WT 2) and 3,437 (*Δ*eMIC 2) cells remained, 12,862 in total. Clustering analysis was performed using the R package *Seurat* (v 4.3.0)^140^ as described in the tutorial (http://satijalab.org/seurat/). In brief, gene expression matrices were SCT-transformed and a default of 3,000 variable genes were selected for batch integration and PC analysis^141^. The top 16 PCs were used for clustering as determined using the *findPC* R package (v 1.0)^142^. Clustering was performed on the integrated data with the Louvain modularity algorithm (*FindClusters* function, resolution = 0.6) to generate 20 biologically meaningful clusters. Assignments of cluster identities were based on candidate marker genes obtained from the *Seurat FindMarkers* function with requirements of > 20% expression in the cluster of interest and a log2FC > 0.25 between cells in the cluster of interest. Statistics by Wilcoxon-rank sum test. Marker genes were compared to databases (*DanioCell*; single-cell atlas) and literature^143–145^. The ribosomal gene expression enriched cluster (Ribo^+^) was excluded from downstream analysis. Differential contribution of WT and *srrm3*^ΔeMIC^ to single-cell clusters was tested for using *propeller* from the *speckle* R package (v 1.4.0)^68^. P-values were FDR-adjusted for multiple testing. For visualization of *srrm3* gene expression Adaptively-thresholded Low Rank Approximation (ALRA) was used allowing for the interpolation of zero values in sparse single-cell matrices. ALRA imputation was performed via the *ALRA* R package (v 0.0.0.9000) default pipeline with automatic selection of the rank parameter *k*^146^. To compute differential gene expression as input for the GSEAs *Seurat findMarkers* function with > 30% expression in the cluster of interest, using a negative binomial model with batch as latent variable was used. The GSEA analysis was performed using the R package *clusterprofiler* (v 4.12.0)^147^ and the GSEA plot was made using the R package *enrichplot* (v 1.24.0).

#### Differential splicing analysis

Alternative splicing analysis was performed using *vast-tools*^122^. In brief, 150 base pair paired-end reads were mapped to the *Danio rerio* genome (GRCz10) using *vast-tools align*, resulting in multiple output files that were subsequently combined to a single table with *vast-tools combine*. Differential splicing analysis was performed using *vast-tools compare* (parameters: *--min_dPSI 15, --min_range 5, --paired, --sp Dre, --print_AS_ev, -- GO*). F-statistic (cut-off: 3.8) was obtained from the vast-tools inclusion table using the R package *betAS* (v 1.2.0)^148^. Neuron-specific exons were obtained from Lopez-Blanch *et al.*^32^. ΔPSI (% change) was computed as *(ΔPSI ΔeMIC/ΔPSI WT)*100*. For cross-species comparisons of splicing profiles^19,21^ orthologous human/mouse exons were obtained from *VastDB*^122^ and differential splicing tables obtained via *vast-tools* as described above. Events were included if they passed coverage filters across both species and if they did not have one-to-many relationships with zebrafish exons. GO enrichment analysis was performed using the R *clusterProfiler package*^147^ with a list of background genes of exons that passed coverage filters from *vast-tools* compare (*--GO*). The protein impact of exon splicing was determined using exon-level information available at *VastDB.* Categories were simplified into 3 groups (alternative UTR, ORF disruption, alternative protein isoform). Categories “non coding” and “in the coding sequence with uncertain impact” were excluded from the analysis.

#### Differential gene expression analysis

Reads were mapped to the *Danio rerio* transcriptome (GRCz10) using *Salmon* (v 1.5.1) with the minimum accepted length for a viable match = 25^149^. Mapping rates were > 60% for all samples with the exception of ΔeMIC-DMSO (run i) with 29% mapping and contamination (human, mouse) thus excluded from downstream analysis (Table S4). The count table with raw counts and TPMs were imported into R using the R package *tximport* (v 1.32.0)^149,150^. Only genes with > 10 counts and TPM > 1 across replicates in at least one group (meaning genotype or if applicable genotype[treatment]) were kept. Differential gene expression analysis on the high-depth 125 bp paired end sequencing data set was performed using *DESeq2* (v 1.44.00) (model ∼ batch + genotype). Differential gene expression analysis of the shallower 50 bp single end sequencing data set was done with the R package *limma* (v 3.60.0) using “Empirical Bayes Statistics for Differential Expression” with a model of ∼ genotype(treatment)^151^. The input count matrix was obtained from *DESeq2* (v 1.44.00) (model ∼ 1) exposed to a variance stabilising transformation (VST)^152^ that was batch corrected using the “Empirical Bayes-moderated adjustment for unwanted covariates” from the *WGCNA* R package (v 1.72.5) retaining genotype(treatment) as covariate^153^. GO term enrichment (all terms [biological processes, molecular function, cellular component]) (set size: 10-200) and GSEA analysis were performed using the R package *clusterprofiler* (v 4.12.0)^147^. GSEA plots were made using the R package *enrichplot* (v 1.24.0). Volcano plots were made using the *EnhancedVolcano* R package (v 1.22.0). The *STRING* protein-protein interactions were obtained using the R package *STRINGdb* (v 2.16.0) with *STRING* (v 12.0)^154^ with a minimum interaction strength of 200. Network plots were visualized using the *igraph* (v 2.1.4) and *ggraph* (v. 2.2.1). Chromosome information was obtained using the R package *biomaRt* (v. 2.60.1).

#### Sequence similarity between srrm3 regulated exon harbouring genes and paralogous genes

Paralog sets were obtained from *Ensembl* biomart (v 112)^155^. The % sequence similarity was obtained for the paralogs of all *srrm3* regulated microexons (length ≤ 51 nt, ΔPSI < - 15) genes (background set) excluding genes harbouring *srrm3*-regulated microexons themselves and the DEGs to test for (target set). The similarity of background versus target genes was compared using a permutation test. The p-value was calculated as *(r + 1)/(n + 1)*, where *r* is the number of permutations more extreme than the observed value and *n* is the total number of permutations (here n = 1000)^130,131^. Note, a single target/background gene could have multiple similarity scores if it was a paralog of multiple *srrm3* regulated microexon harbouring genes.

### *In situ* hybridization chain reaction (HCR)

#### Probe design

Probes were either purchased from *Molecular Instruments* (*elavl3*), together with the HCR amplifiers (B1-Alexa Fluor 647, B2-Alexa Fluor 546) and required buffers or purchased from *Integrated DNA Technologies* (*fosab*, *srrm3*) following a custom design described in Choi *et al.*^156^. The following combinations were used: B2-*elavl3*, B1-*fosab*, B1-*srrm3*. Any probe pair that fell below a % GC - or melting temperature threshold was excluded, as well as probe pairs with possible off-target effects. Probe pairs provided in Table S5.

#### Experimental design

Exp 1 - *srrm3* expression: *srrm3* staining of 6 dpf WT larvae (N = 1 clutch).

Exp 2 - Brain morphology: *elavl3* staining of 6 dpf *srrm3*^ΔeMIC^ and WT larvae (N = 3 clutches).

Exp 3 - Brain morphology and *fosab* expression: *elavl3* and *fosab* staining of 6 dpf *srrm3*^ΔeMIC^ and WT larvae (N = 3 clutches).

#### Fixation

Larvae were fixed at 6 dpf during the day with the exception of Exp 3 (*elavl3, fosab*) for which larvae were fixed at night (2 h after light off). Petri dish contents were poured directly into a 40 µm nylon cell strainer (Cultek - 88141378), briefly dried to remove excess media, and immediately immersed in 4 % freshly prepared paraformaldehyde (PFA) in 1 x PBS. Larvae were transferred into 1.5 ml Eppendorf tubes using a pasteur pipette and either fixed at 4 C overnight (ON) or at room temperature (RT) for 3 h while on a rotorad. Larvae were either fin-clipped for genotyping at 3 dpf^121^ or after fixation in 1x PBS, before step-wise dehydration into 100% MetOH final for -20 storage (25 %, 50 %, 75 % in 1 x PBS). *srrm3*^ΔeMIC^ and WT larvae were stained in different tubes.

#### HCR: Experimental procedure

The HCR staining was performed on larvae with a homozygous *nacre* background (*mitfaw2* allele) to avoid skin pigmentation^60^ as described by Bruce *et al.*^157^ adapted from Choi *et al.*^156^. Up to 21 larvae were stained in the same tube. In brief, larvae were stepwise dehydrated from MeOH and then washed 3 x in PTw (1 x PBS, 0.1 % Tween 20). Larvae were permeabilized and then pre-hybridized in probe hybridization buffer (*Molecular Instruments*) at 37 C for 30 min. This pre-hybridization solution was removed and a probe hybridization buffer with 0.8 pmol of probes was added for incubation ON at 37 C. Larvae were washed in the probe wash buffer (*Molecular Instruments*), followed by washes with 5 x SCCT (sodium chloride sodium citrate, 0.1% Tween 20). Samples were then pre-amplified in 1 ml of amplification buffer (*Molecular Instruments*) for 30 min at RT. Meanwhile 2 µl (3 µM stock) of each hairpin h1 and h2 of each amplifier (B1, B2) was added to the amplification buffer and heated for 90 sec at 95 C before cooled down to RT in the dark. The pre-amplification solution was removed, the hairpin solution added and the larvae incubated in the dark ON at RT. Larvae were repeatedly washed in 5x SCCT and transferred to 1 x PBS. DAPI was added as a nuclear stain for ∼ 1h at 1.0µg/mL. Samples were stored at 4 C in 1 x PBS for up to 4 days before imaging.

#### Mounting and Imaging

Around 5 - 8 larvae were mounted onto each glass bottom MatTek dish (35 mm; P35G-1.5-20-C) using 1.5% low-melting point agarose (LMA; SeaPlaque™ Agarose [50100]) in 1 x PBS oriented dorsal down. Imaging was done with an Axio Observer.Z1/7 Zeiss LSM 980 microscope acquiring imaging stacks with the EC Plan Neofluar 10x/0.30 M27. The scan zoom was set at 1.2 x zoom with bidirectional imaging. Imaging was performed using a GaAsP-PMT detector. Stacks were collected at a z-spacing of 4 µm with the exception of *srrm3* stainings which were collected at 5 µm. Exp 1 (*srrm3*): The laser wavelength was 639 nm (4%, detector gain: 770V) for B1-Alexa Fluor 647. Exp 2 (*elavl3*): The laser wavelength was 561 nm (0.9%, detector gain: 700V) for B2-Alexa Fluor 546. Exp 3 (*fosab*, *elavl3*): The laser wavelength was 639 nm (3%, detector gain: 750V) for B1-Alexa Fluor 647 nm and 561 nm (2.2%, detector gain: 700V) for B2-Alexa Fluor 546.

#### Image registration and visualization of srrm3-labelled brains (Exp 1)

3D brain registration was performed using the ANTs toolbox (v2.3.5)^136^ to the ZBB reference^134,135^, following the approach described in Antinucci *et al.*^132^. *srrm3* stainings are shown as maximum z-projection of sagittal and dorsal views generated with *Fiji/ImageJ2* (v 2.14.0/1.54f). Using a mask of the whole ZBB brain, areas outside of the brain were set to zero. The colour scale was adjusted (Fire).

#### Morphological analysis (Exp 2,3)

*elavl3*-labeled brains for anatomical reference were registered to an in-house reference brain using the Computational Morphometry Toolkit (CMTK)^158–161^. The MapMAPPING pipeline^160^ was used for the analysis of brain volume changes as previously described^69,161^. The FDR threshold was set to the default of 0.00005.

#### Quantification and visualization of fosab stainings (Exp 3)

3D brain registration was performed using the ANTs toolbox (v2.3.5)^136^ to the ZBB reference^134,135^, following the approach described in Antinucci *et al.*^132^. In *MATLAB*, following registration, the difference in background intensity was addressed by subtracting the mode of pixel intensities from the image stacks, providing a reliable estimate of background or baseline fluorescence, similarly to Tunbak *et al.*^162^. As a control strategy we also performed multi-otsu thresholding in *Python* from the *scikit-image* package^163^ to distinguish between background and signal which resulted in more stringent thresholding^124^ but led to similar results in the downstream analysis. Pixel-wise comparisons of *fosab* signal intensity across brains and intensity maps were obtained using the MapMAPPING pipeline^160^ with input tiff stacks at 8 bit, downsampled to 300 x 679 x 80 (x, y, z) and pre-smoothed with a 2D gaussian filter (sigma = 1). The FDR threshold was set to the default of 0.00005. Stacks of significant delta medians were visualized with *Fiji/ImageJ2* (v 2.14.0/1.54f) by obtaining maximum intensity projections of coronal and sagittal stacks overlaid with the Z-brain atlas^160^ outline.

### Pharmacological assays

#### Behavioural pharmacology

Sleep/wake experiments were performed as described in “*Sleep/wake experiments and analysis*”. Larvae were plated the evening before experimental day 1 (5 dpf) into alternating columns of wells with either drug or control in E3 media that were pre-mixed in 50 ml falcon tubes. Larvae were moved with a P200 that was cut at the tip. A gentle tap with the pipette tip against the well edge resulted in the larva entering the well without transferring additional media. The drug/DMSO-E3 mix was used for the water refill every morning. For ropinirole and clonidine the percentage change in activity (% time active) was calculated following treatment compared to the same GT control group: *([activity - activity mean of same GT control]/|activity mean of same GT control|)*100*. A nonlinear dose-response model was fitted to the percentage changes using log10-transformed concentrations and a four-parameter logistic function (Hill equation) with the lower asymptote fixed at zero using *drm* from the *drc* R package (v 3.0-1). For SQ22536, forskolin and rolipram (rescue and mimic conditions) the percentage change in activity (% time active) was calculated compared to the WT control: *([activity - activity mean of WT control]/|activity mean of WT control|)*100.* Statistics via Wilcoxon-test.

Rolipram was purchased at 10 mM in 100% DMSO and subsequently diluted to a stock concentration of 1 mM in 10% DMSO in H2O (rolipram: HY-16900, MedChemExpress). Ropinirole and forskolin were ordered as powder and similarly diluted to a stock concentration, first of 10mM in 100% DMSO and then further to 1mM in 10% DMSO in H2O (ropinirole hydrochloride: R2530-100MG, Sigma; forskolin: F6886- 10MG, Sigma). For each experiment the % DMSO in the control concentration was kept equal to that of the treatment condition. For example, 5µM of rolipram from a 10% DMSO stock would result in 0.05% of DMSO. The same amount of DMSO (0.05%) was also added to the controls. The maximum % DMSO used for any experiment was 0.2% for 20µM of ropinirole treatment. Clonidine was ordered as powder and diluted first to 10mM in H2O and further to 1mM in H2O (Clonidine hydrochloride: C7897-25MG, Sigma). SQ22536 was ordered as powder and diluted to 10mM in 1 x PBS (SQ22536: HY-100396, [25 mg], MedChemExpress). Similarly to DMSO, the control and treatment conditions were exposed to equal amounts of H2O or 1 x PBS. The range of concentrations tested were chosen based on the literature, starting with 2.5µM for clonidine and ropinirole^75^, with 5µM for forskolin, rolipram^43^ and SQ22536. For SQ22536 in particular no apparent behavioural changes were observed at concentrations that resulted in behavioural effects for clonidine, ropinirole, forskolin or rolipram, prompting us to increase concentrations up to 500µM^164^.

#### RNA-seq of pharmacologically treated larvae

Samples were prepared, FACS-sorted and sequenced as described in “*Bulk and single-cell RNA-seq*”. Larvae were exposed to treatment + DMSO or to a matched % of control DMSO (maximum 0.1%) the afternoon prior to dissociation (Table S4), matching the timing that treatments were added when larvae were tracked for behavioural experiments. Concentrations were chosen based on behavioural data to either reduce ΔeMIC daytime activity to that of WT larvae (clonidine: 10µM; ropinirole: 10µM, SQ22536: 500µM) or to increase WT daytime activity to that ot ΔeMIC larvae (rolipram: 0.75µM, forskolin: 7.5µM). PTZ treatment concentration (1mM) was chosen to increase neuronal activity without inducing seizures^165^.

#### Survival assay

Zebrafish larvae (4 - 6 dpf) were exposed to either the mitochondrial calcium uniporter inhibitor ruthenium red at 1.25µM (R2751-1G, Sigma; stock = 1mM in H2O) or 4.7µM of Ru265 (SML2991-5MG, Sigma; stock = 2mg/ml in H2O) shown to target the mitochondrial calcium uniporter at higher specificity^166^. The treatments were added to the petri dish with larvae and E3 (no methylene blue) in the morning and the number of dead larvae was counted every hour from exposure. Larvae of all genotypes were kept in the same petri dish (40 ml total) to be exposed to the same concentration of the drug. We monitored survival hourly for 11 - 14 h. Larvae still alive at the last instant of measurement were considered in the Kaplan-Meier survival method. Larvae were genotyped post-experiment^32^, hence the experimenter was blinded to the larva’s genotype. Statistics were performed in R using a Pairwise comparisons with Log-Rank test from the *survminer* package (v 0.5.0).

### ATAC-sequencing

#### Sample preparation and sequencing

ATAC-seq was performed following FACS-sorting of 60,000 GFP-positive neurons [Tg(*elavl3*:GFP)] per sample from 2 biological replicates of enucleated 6 dpf larvae (N = 2, clutch 1: n = 11 larvae each, clutch 2: n = 12 larvae each). Cells were spun at 120 g for 5 min and the supernatant was removed. For nuclei preparation, the cell pellet was resuspended in 600 µl of nuclear extraction buffer (250 mM sucrose, 25 mM KCl, 5 mM MgCl2, 20 mM HEPES-KOH [pH 7.8], 0.5% IGEPAL CA-630, 0.2 mM spermidine, 0.5 mM spermidine, nuclease-free water) and homogenized on ice using a Dounce homogenizer (Sigma) following Fernandez-Albert *et al.*^88^. The nuclei were spun at 1000 g for 7 min, and the supernatant removed carefully followed by transposition and purification as described in Buenrostro *et al.*^167^. Briefly, the pellet was resuspended in 50 µl transposase reaction mix (25 µl 2x TD buffer, 3.5 µl TN5 [ref. 20034210, Nextera, Illumina], 21.5 µl of nuclease-free water) and transposed for 30 min at 37 C and 400 rpm, followed by purification with a MinElute PCR purification kit (ref. 28004, Qiagen). At the CRG Genomics facility: Library amplification and barcoding were performed with NEBNext Q5 Hot Start HiFi PCR Master Mix (ref. M0543L, New England Biolabs) using dual index nextera primers, at a final concentration of 1.25 µM. PCR was conducted for 12 - 14 cycles. Library purification was performed with AgenCourt AMPure XP beads (ref. A63882, Beckman Coulter). Final libraries were analyzed using Agilent Bioanalyzer or Fragment analyzer High Sensitivity assay (ref. 5067-4626 or ref. DNF-474) to estimate the quantity and check size distribution, and were then quantified by qPCR using the KAPA Library Quantification Kit KK4835 (ref. 07960204001, Roche). Libraries were sequenced 2 * 51+10+10 base pairs on Illumina’s NovaSeq6000 with 67-70 million reads per sample (clutch 1: WT1 + ΔeMIC1; clutch 2: WT2 + ΔeMIC2) (Table S4).

#### ATAC-sequencing data processing and analysis

Read quality control and adapter trimming were performed using *trimmomatic* (v 0.39)^168^ [SLIDINGWINDOW:4:20; ILLUMINACLIP:AT_Tn5_adapters_NexteraPE-PE.fasta:2:30:10]. Reads were mapped to the danRer10 genome using *bwa* (v 0.7.18-r1243-dirty)^169^, deduplicated and coordinate-sorted using *bamsormadup* from *biobambam2* (v 2.0.183) in combination with *samtools* (v 1.21)^170^, resulting in ∼ 50 million paired-end reads per sample. Next *alignmentSieve* from *deeptools* (v 3.5.5)^171^ was used for read shifting to the transposon binding event^172^ and filtering of nucleosome-free fragments (1 - 120 base pairs). Peak calling was performed on nucleosome-free fragment merged .bam files from all samples, obtaining consensus peaks using *macs3* (v 3.0.2)^173^ (narrow mode, nomodel, q = 0.01). For the count table generation with *featureCounts* (v 2.0.8)^174^ all fragment lengths were included with reads requiring 20% minimum overlap, resulting in ∼ 13 million successfully assigned alignments per sample. Peaks were annotated using *annoPeaks* from the R package *ChIPpeakAnno* (v 3.38.1)^175^. Differential footprinting scores were computed only on samples from clutch 1 due to a shifted fragment length distribution of ΔeMIC2 (fragment length medians: WT 1 = 103; ΔeMIC 1 = 103; WT 2 = 104; ΔeMIC 2 = 262) using *TOBIAS*^176^ with protein family database (*PFAM*) motifs from *JASPAR* in *MEME* format (750 non-redundant vertebrate motifs)^177^.

## Author contributions

Conceptualization, TM, MI; Methodology, LPI, TSM, FK, GZ, JFA, JP, IHB, MO, JR, MI; Software, TM, LPI, TSM, FK, GZ, IHB, MO, JR, MI; Validation, TM, CRM, LSV; Formal Analysis, TM, LPI; Investigation, TM, TSM, CRM, GZ, JFA, LSV, JP; Resources, TM, CRM, IHB, MO, JR, MI; Writing - Original Draft, TM, MI; Writing - Review & Editing, TM, TSM, FK, JFA, IHB, MO, JR, MI; Visualization, TM, MI; Supervision, TM, IHB, MO, JR, MI; Project Administration, MI; Funding Acquisition, IHB, MO, JR, MI.

## Acknowledgements

We thank Federica Mantica, Ludovica Ciampi, Sophie Bonnal, Jonas Juan-Mateu, John Chamberlin, Antonio J. Montero-Hidalgo, Niccolo Arecco for their critical reading of the manuscript and feedback. Snigdha Nadagouda and Elena Putti are gratefully acknowledged for their help on developing strategies of experimental validation and Chintan Trivedi for designing the HCR probe sequences. We also thank the Sebé-Pedrós lab for valuable advice on sequencing experiments. A great thank you to present and past members from the TransDevo lab and the UCL fish floor for their invaluable support and feedback throughout. We also thank the CRG Genomics Unit for assistance with the sequencing, the CRG/UPF Flow Cytometry Unit for assistance with the FACS, and the CRG Advance Light Microscopy Unit for assistance with the confocal image acquisition. The research has been mainly funded by the European Union’s Horizon 2020 research and innovation program through the European Research Council (ERCCoG-LS2-101002275 to MI) and the Marie Skłodowska-Curie MSCA-ITN-ETN scheme (project ZENITH, grant agreement No 813457 to MI, IHB and MO, which provided funds for TM, TSM and GZ). The research has also received funding from the Spanish Ministry of Science and Innovation (PID2020- 115040GB-I00 to MI), a Wellcome Trust Investigator Award (#217150/Z/19/Z to JR), a BBSRC Research Grant (BB/X01536X/1 to JR), a European Research Council (ERC NEUROFISH 773012 to MO), Volkswagen Stiftung ”Life?” initiative (to MO). LPI acknowledges support from IJC2020-044783-I funded by MCIN/AEI/10.13039/501100011033 and by European Union NextGenerationEU/PRTR, and JFA a Beatriu de Pinós fellowship (2021 BP 00195). CRG acknowledges support of the Spanish Ministry of Science and Innovation through the Centro de Excelencia Severo Ochoa (CEX2020-001049-S, MCIN/AEI/10.13039/501100011033), and the Generalitat de Catalunya through the CERCA program.

## Supplementary Table legends

**Supplementary table 1: Statistics and sample sizes of pixel-based behavioural assays.** Model estimates ± SE and sample sizes for the thigmotaxis assay, the dark-light assay, the tapping habituation assay and all sleep/wake parameters tested^124^ for replicates of *srrm3* the *vsx1;vsx2* experiment and the *srrm3;srrm4* (Fig. 1) but also for the individual microexon mutants (Fig. 4J).

**Supplementary table 2: Statistics and sample sizes for the bout type and bout kinematics analysis.** Sample sizes and statistics for bout usage and kinematics, long bouts and PCA/Euclidean distance, specifying when individual bouts or larvae were excluded from the analysis (Fig. 2).

**Supplementary table 3: Footprinting output using *TOBIAS*.** Statistics, including footprinting scores and number of transcription factor binding sites (TFBSs) for each of the tested motifs, obtained as output from *TOBIAS*^176^. The TFBSs associated with each motif are shown with information on the peak annotation and gene (ENSEMBL ID) added.

**Supplementary table 4: Overview of sequencing experiments.** Information about bulk (Fig. 4, Fig. 6, Fig. 7), single-cell (Fig. 4, Fig. 5), and ATAC-sequencing experiments (Fig. 7), including sample size, treatments, RNA integrity number (RIN), life cells (%) after FACS-sorting, paired vs single end, sequencing depth, read length, mappability, the type of analysis the data was used for.

**Supplementary table 5: HCR probes.** Sequences of the oligonucleotides ordered for the HCR probes of *fosab* and *srrm3*.

## Supplementary figures

**Supplementary figure 1:**
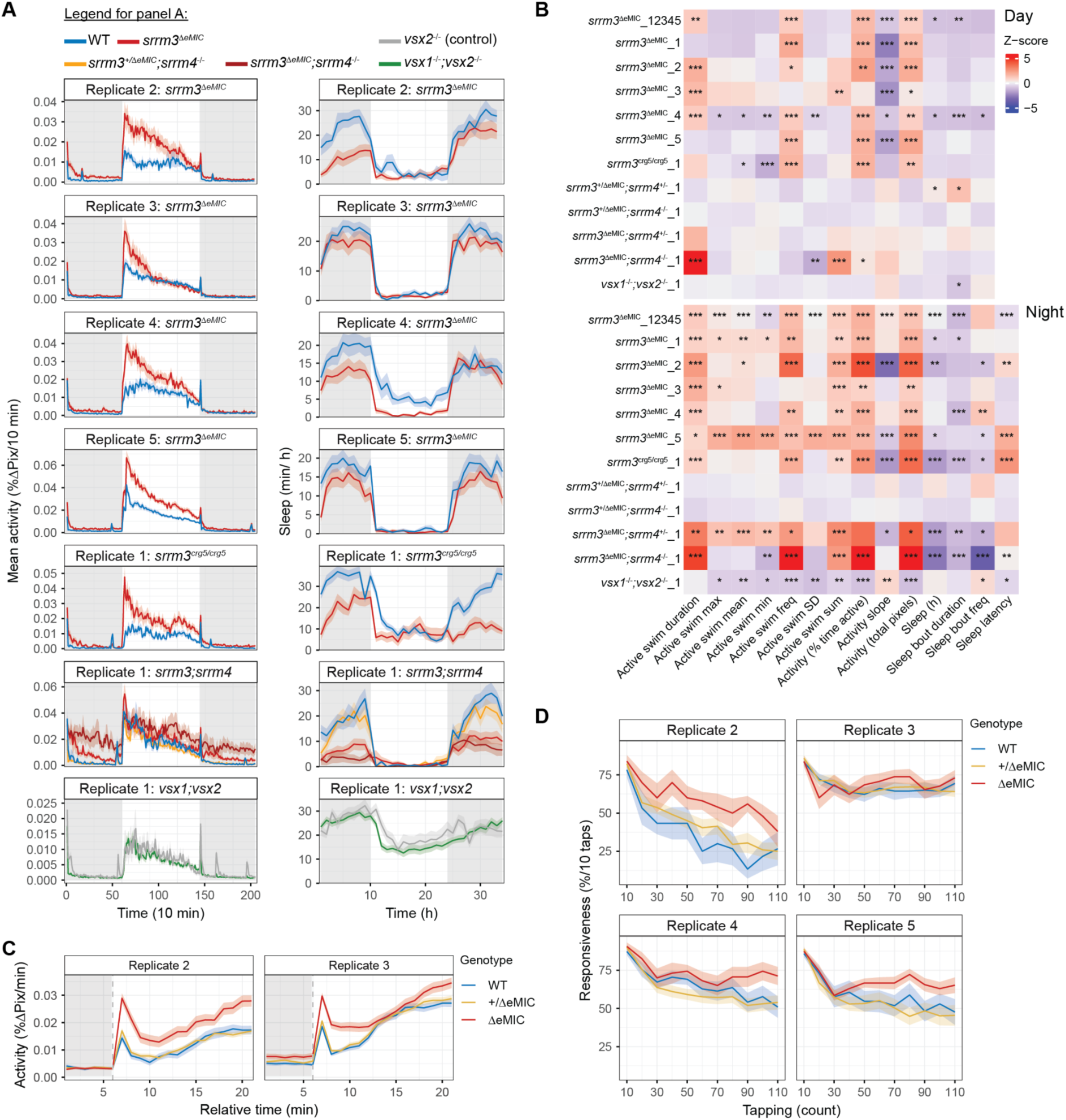
*srrm3*^ΔeMIC^ larvae show hyperactivity, sleep loss, sensory hypersensitivity and excess thigmotaxis. A: Representative traces (mean ± SEM) for activity (%ΔPix/10 min) (left) and sleep (min/h) (right) during 74 h (5 - 8 dpf) on a 14 h light - 10 h dark cycle (white background and grey background respectively). For visualisation, values of refill periods were replaced with pre refill values. For clarity heterozygous traces are not shown (except for the double mutant) as they resemble WT traces. B: Summary of behavioural parameters analysed^124,178^ during the day (6 dpf) and at night (5 - 6 dpf and 6 - 7 dpf). Average z-scores across larvae standardized to the respective WT or control [*vsx2*^-/-^] sibling. Number after the gene name (1 to 5) indicates biological replicate or averaged replicates in the case of 12345 for which LMM statistics include all 5 clutches. Stars indicate significance as tested by likelihood-ratio test on LMM and represent: *p < 0.05, **p < 0.01, *** p < 0.001. C: Traces (mean ± SEM) of averaged (5 transitions per larvae) light-on response at 6 dpf of activity (%ΔPix/min) across larvae from the same clutch and genotype. N = 2 clutches with 12 to 45 larvae each per genotype. D: Traces (mean ± SEM) of habituation to mechanical tapping stimuli at 6 dpf showing mean responsiveness (%) to 10 tapping stimuli at 90 sec inter-stimulus interval (tap 1- 10) and 5 sec inter-stimulus interval (11 - 100). N = 4 clutches with 5 to 46 larvae each per genotype. A,C,D: Biological replicates are indicated counting from 2 since replicate 1 is shown in the main figure. B-D: Detailed statistics and sample sizes in Table S1.

**Supplementary figure 2:**
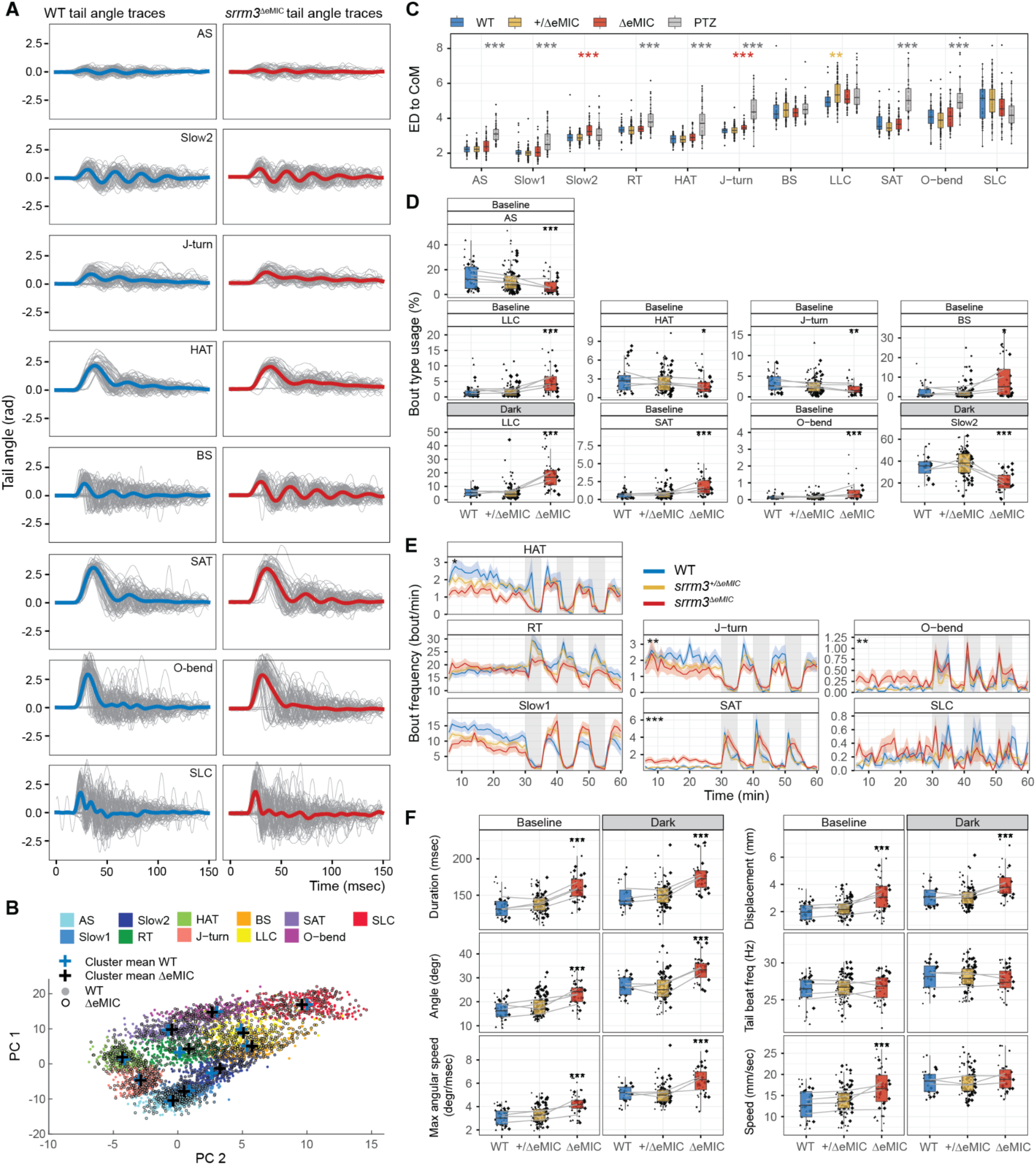
Bout type usage and kinematics reflects a heightened arousal state in *srrm3*^ΔeMIC^ larvae. A: Representative tail traces (7^th^ tail segment) across 8 bout types (the remaining 3 are shown in Fig. 2B) with 70 randomly sampled individual traces per type (grey) and mean trace shown in bold, from one clutch with 8 WT and 14 *srrm3*^ΔeMIC^ larvae. B: Bouts clustering in PC space of kinematics^50^. Crosses indicate cluster means by genotype for WT (blue, n = 41 larvae) and *srrm3*^ΔeMIC^ (black, n = 46 larvae). For each bout type 200 bouts were randomly sampled. C: Median Euclidean distance (ED) per larvae (dot) and bout type to the cluster centre of mass (CoM)^50^. Statistics by Wilcoxon-test with stars coloured by groups compared to WT. P-values FDR corrected for testing across multiple bout types. Y-limits for visualisation: min = 1.5, max = 8.5. Sample size: n = 41 WT, 110 *srrm3*^+/ΔeMIC^, 46 *srrm3*^ΔeMIC^, 43 10mM PTZ-treated larvae of mixed genotype. D: Bout type usage (%) for types significantly different between WT and *srrm3*^ΔeMIC^ siblings, split by condition (baseline, dark). E: Mean bout frequency (bout/min) over time, with grey background indicating dark periods. Traces represent mean ± SEM across larvae for each genotype. Significant stars are added for comparisons during baseline and darkness. D,E: Statistics by maximum likelihood test on LMM on log-transformed values. P-values were Bonferroni corrected for testing multiple bout types in each condition (baseline, dark). N baseline = 4 clutches, dark = 3 clutches, with 7 to 33 larvae each per genotype. F: Mean values across bout kinematic parameters split by bout type and condition (baseline, dark). Statistics by maximum likelihood test on LMM with p-values Bonferroni corrected for the number of bout types tested for each condition. N baseline = 4 clutches, dark = 3 clutches, with 7 to 33 larvae each per genotype. D,F: Each dot represents one larva shaped by clutch. Grey lines connect clutch means. C-F: Significance stars represent: *p < 0.05, **p < 0.01, *** p < 0.001. Statistics shown between WT and *srrm3*^ΔeMIC^ siblings with details and exact sample sizes in Table S2. A-E: AS, approach swim; RT, routine turn; HAT, high-angle turn; BS, burst swim; SAT, spot avoidance turns; SLC, short-latency C-starts; LLC, long-latency C-start.

**Supplementary figure 3:**
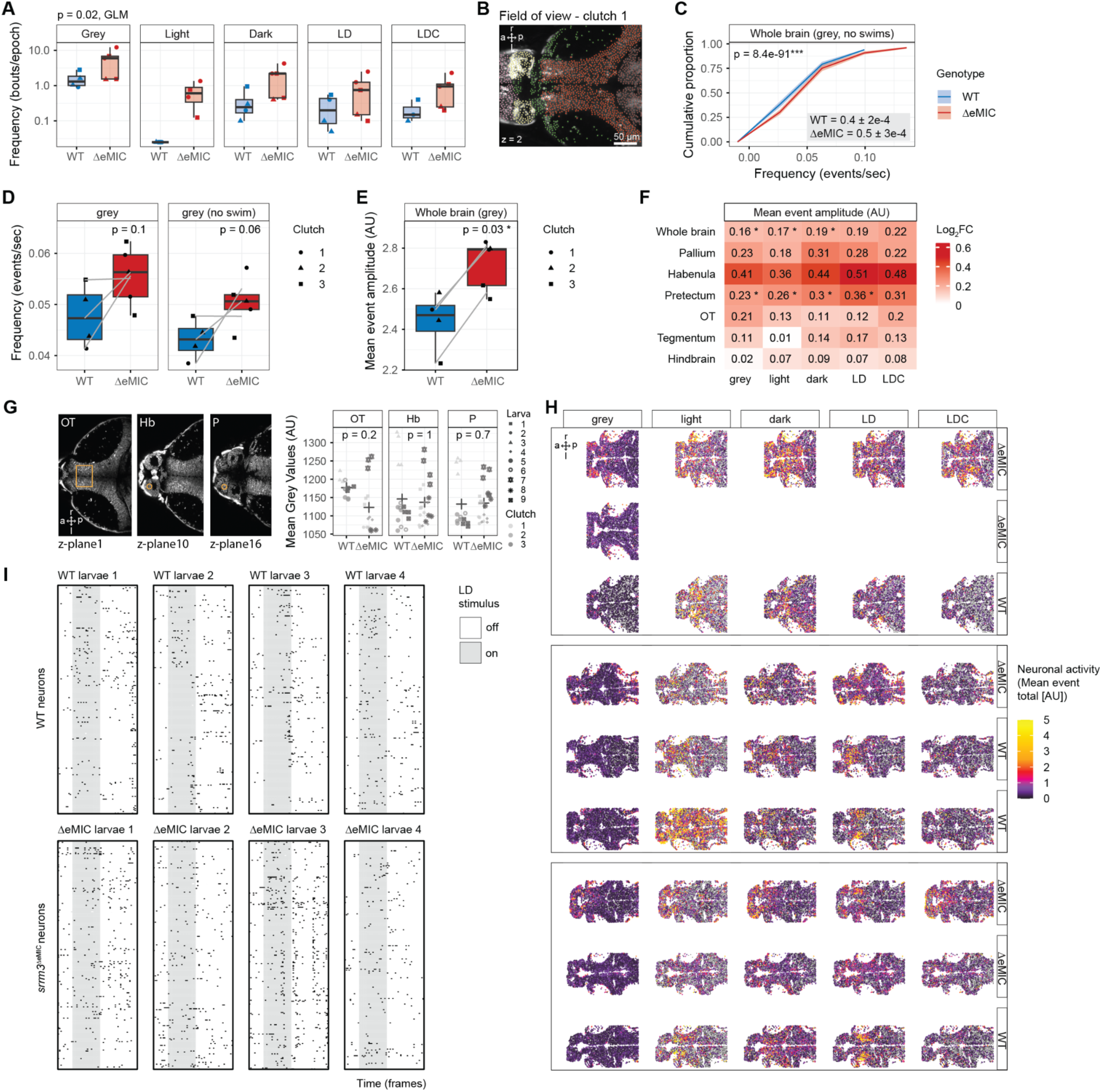
*srrm3*^ΔeMIC^ larvae show increased neuronal activity at baseline and a distinct response to visual stimuli. A: Swim bout frequency (bouts/epoch) across stimuli on log10- transformed y-axis (frequency: 1.3 ± 0.6 bouts/epoch, p = 0.02). Each dot represents one larva shaped by clutch. Statistics by GLM controlling for stimulus and larva (n = 4 WT and 5 *srrm3*^ΔeMIC^ larvae from 3 clutches). B: FOV for clutch 1 showing one z-plane with ROIs coloured by ZBB anatomical region. C: ECDFs of event frequency for the whole brain at baseline, excluding periods of swimming. Lines indicate means ± SEM across the same genotype and are cut at y = 0.995 for visualization. Boxed values show mean ± SEM across neurons of the same genotype. Statistics by Wilcoxon-test, comparing 36,951 WT and 38,940 *srrm3*^ΔeMIC^ neurons from 4 WT and 5 *srrm3*^ΔeMIC^ larvae. D: Event frequency across the whole brain at baseline considering all frames (left) and excluding frames with swims (right). E: Mean event amplitude across the whole brain at baseline. D,E: Dots represent larva means. Lines connect means across clutches. Statistics by Wilcoxon-test, on n = 4 WT and 5 *srrm3*^ΔeMIC^ larvae from 3 clutches. F: Heatmap of event amplitude log2FC. Blue indicates higher amplitudes in the WT and red in *srrm3*^ΔeMIC^. Statistics by Wilcoxon-test, comparing 4 to 5 larvae per genotype from 3 clutches. Significance stars represent: *p < 0.05, **p < 0.01, *** p < 0.001. G: Measurements of *Gcamp6s* fluorescence (mean intensity ) at 800 nm with laser power and gain settings identical across animals. Representative larva anatomies showing ROIs (orange outline) used for measurements in the optic tectum (OT), habenula (Hb) and pallium (P). Statistics by Wilcoxon-test on mean values per larva and region. n = 4 WT and 5 *srrm3*^ΔeMIC^ larvae with 3 to 4 measurements per brain region from 3 clutches. H: Anatomy plots of one z-plane of all larvae (rows) and stimuli (columns) with neurons (dots) coloured by mean event total (AU). Values > 5 were set to 5 for visualisation. Boxes indicate larvae from the same clutch. I: Events (black dots) of 243 habenula neurons (columns) of across n = 4 larvae per genotype over time (rows) for 60 frames (∼ 15 sec) with the looming dot stimulus window in grey. For each larva the first well-aligned LD epoch was chosen. B,G,H: a, anterior; p, posterior; r, rightwards; l, leftwards.

**Supplementary figure 4:**
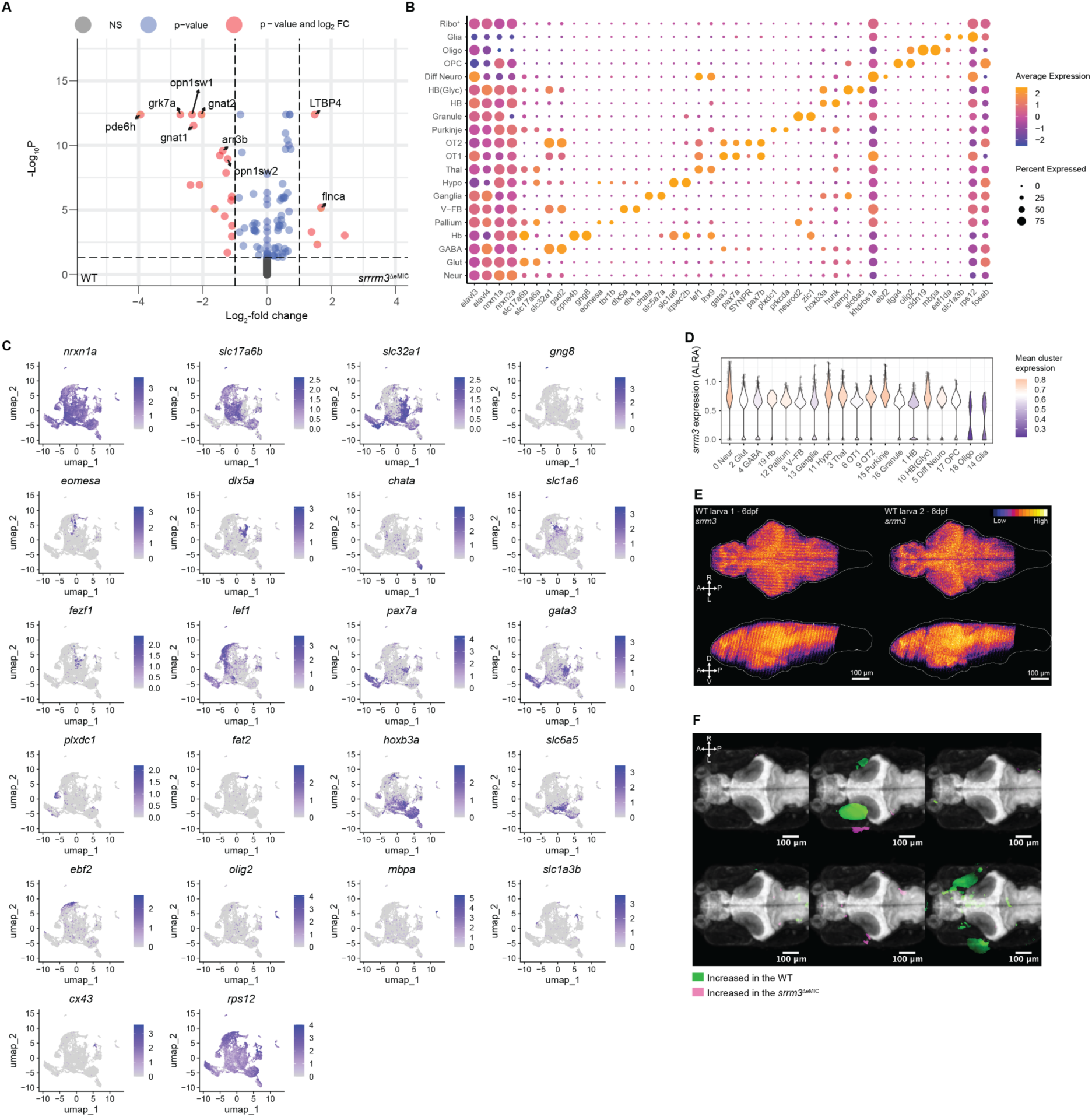
*srrm3*^ΔeMIC^ larvae show reduced microexon inclusion and no differences in cell type composition. A: Volcano plot showing the 8000 most differentially expressed genes (DEGs; ranked by log2FC) in *srrm3*^ΔeMIC^ compared to WT neurons of non-enucleated 5 dpf larvae. Cut- offs for visualisation: p.adj < 0.05, |log2FC| > 1. Genes upregulated in *srrm3*^ΔeMIC^ on the right and downregulated on the left. N = 2 biological replicates per genotype with 20-26 larvae per replicate. B: Two out of the top ten candidate marker genes (columns) for each cluster (rows) are shown. Markers were expressed in > 20% of the cluster and with a mean log2FC > 0.25 compared to all other clusters. Statistics by Wilcoxon-rank sum test. N = 2 biological replicates per genotype with 13-21 larvae per replicate. C: UMAPs of marker genes, with each dot coloured gene expression (log-transformed and corrected counts). Neuronal: *nrxn1a^+^*; GABAergic: *slc32a1^+^*; Glutamatergic: *slc17a6b^+^*; Habenula: *gng8^+^*; Pallium: *eomesa^+^*; Subpallium/hypothalamus/preoptic: *dlx5a^+^*; Ganglia (acetylcholine^+^): *chata^+^*; Hypothalamus: *slc1a6*^+^ (glutamatergic), *fezf1*^+^ (GABAergic); Dorsal diencephalon (thalamus): lef1^+^; OT1: *pax7a^+^*; OT2: *gata3^+^*; Purkinje: *plxdc1^+^*; Cerebellum (granule): *fat2^+^*; Hindbrain: *hoxb3a^+^*; Hindbrain (glycinergic): *slc6a5^+^*; Differentiating midbrain: *ebf2^+^*; Oligodendrocytes: *olig2^+^*; OPCs: *mbpa^+^*; Glia: *slc1a3b^+^*, *cx43^+^*; Ribosomal: *rps12^+^*. D: *srrm3* expression across WT clusters colored by relative cluster average (purple = below, white = average, orange = above). Expression is log-transformed and corrected counts with ALRA used for imputation on sparse single-cell data^146^. Dots represent single cells. E: Maximum intensity z-projections of two WT larva zebrafish brains (larva 1, 2) HCR-stained for *srrm3* transcript dorsally (top) and sagittally (bottom). Brighter colours indicate higher expression levels. F: Brain structural phenotypes obtained from MAPMapping^160,161^ overlaid on the maximum intensity z-projection of an *elavl3*-stained in-house reference brain. Colours indicate significant delta means. Sample size: N = 6 clutches with 8 to 11 larvae per genotype and comparison. E,F: A, anterior; P, posterior; R, rightwards; L, leftwards; D, dorsal; V, ventral. B,D: Cluster names as in Fig. 4.

**Supplementary figure 5.**
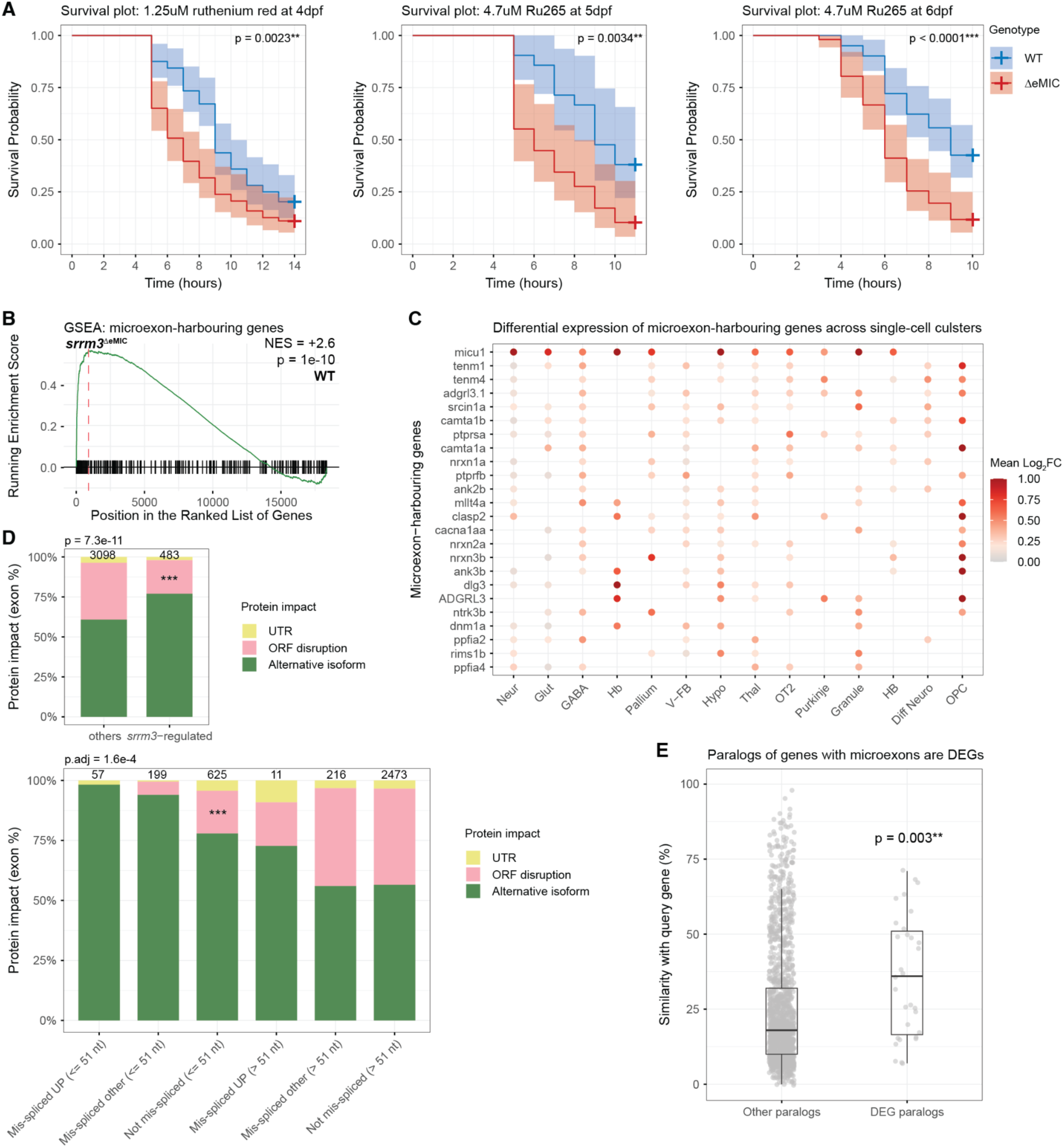
Genes upregulated in the *srrm3*^ΔeMIC^ disproportionally harbour mis- spliced microexons. A: Kaplan-Meier plots showing survival probabilities over time (h post treatment) for mitochondrial calcium uniporter inhibitors: 4 dpf 1.25 µM ruthenium red-treated larvae (left: N = 4 clutches with 11 to 19 larvae each per genotype), for 5 dpf 4.7 µM Ru265-treated larvae (2.5 µg/ml) (centre, N = 2 clutches with 6 to 30 larvae each per genotype) and for 6 dpf 4.7 µM Ru265-treated larvae (right: N = 4 clutches with 8 to 20 larvae each per genotype). Traces represent survival probability ± 95% confidence intervals across larvae from the same clutch and genotype. Statistics by log-rank (Mantel-Cox) test. B: GSEA on log2FC between WT and *srrm3*^ΔeMIC^ for microexon-harbouring genes (length ≤ 51 nt, ΔPSI < -15). C: *srrm3*-regulated microexon-harbouring genes (rows) contributing to the GSEA core enrichment in > 5 clusters (columns), coloured by averaged log2FC. For visualisation maximum fold-changes were set to 1. Cluster names as in Fig. 4. D: Protein impact of a given exon, either disrupting the open reading frame (ORF), resulting in an alternative protein isoform or an alternative untranslated region (UTR). Exon number is indicated at the top. Statistics via two-sided Fisher exact tests comparing the number of exons disrupting the ORF in each group. Left: *srrm3*- regulated (|ΔPSI| > 15) vs. other exons (odds ratio = 0.5, p = 7.3e-11). Right: Groups split by mis-spliced (|ΔdPSI| > 15), UP (upregulation of the exon-harbouring transcript) and exon length (≤ 51 nt or > 51 nt). Comparing mis-spliced exons of upregulated genes with same-size groups revealed statistical significance only for mis-spliced UP (≤ 51 nt) vs. mis-spliced other (≤ 51 nt) (odds ratio = Inf, p.adj = 1.6e-4). P-values were Bonferroni adjusted. E: Differentially expressed paralogous genes show a higher percentage similarity to genes harbouring *srrm3*-regulated microexons (length ≤ 51 nt, ΔPSI < - 15) (query) than all other paralogs. P-value corresponds to a Permutation test. Each dot represents one paralog pair.

**Supplementary figure 6:**
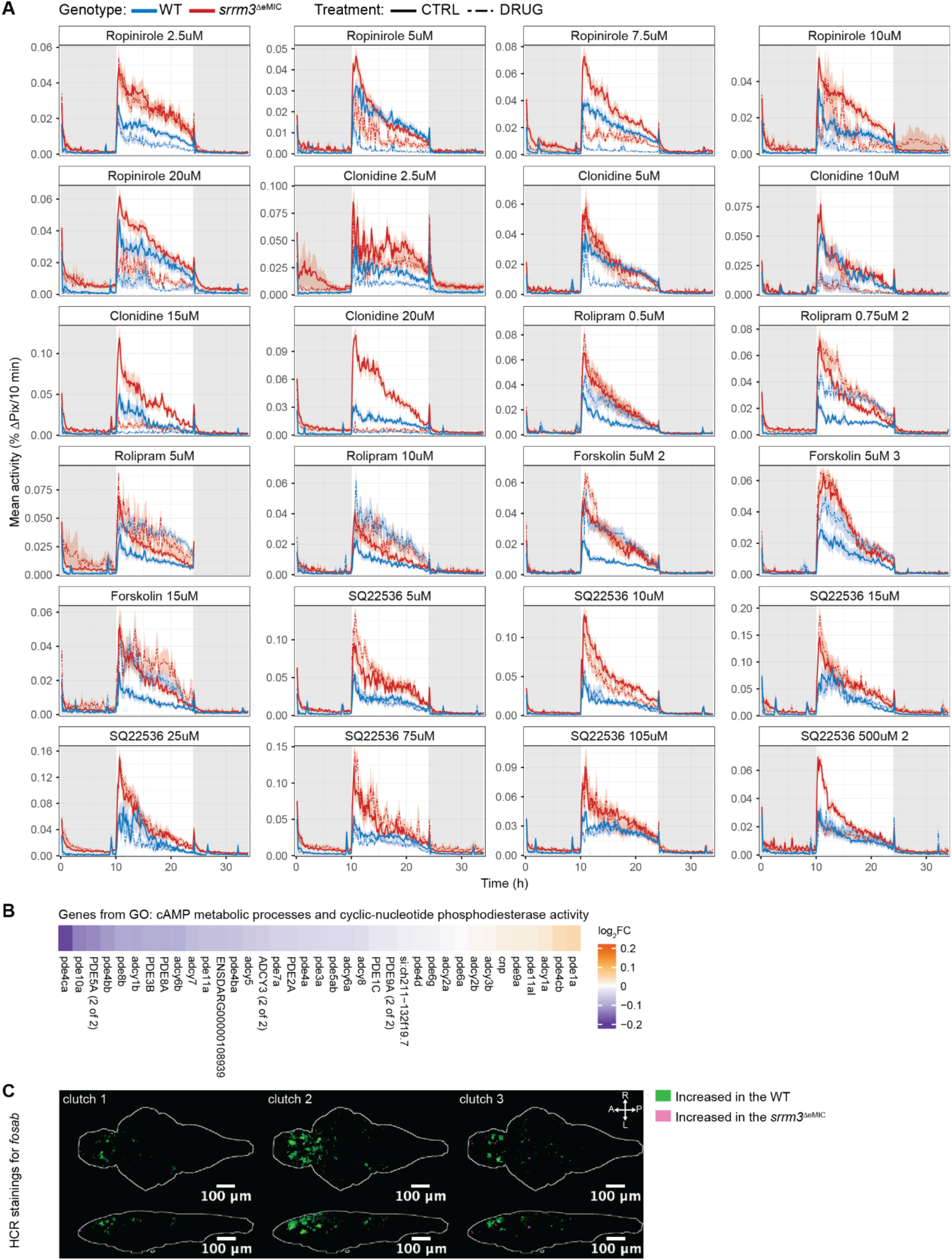
cAMP signaling is central to *srrm3*^ΔeMIC^ daytime hyperactivity. A: Mean activity (%ΔPix/10 min) shown for *srrm3*^ΔeMIC^ (red) and WT siblings (blue), either drug-treated (dashed line) or control (solid line) during 34 h (5 - 7 dpf) on a 14 h light - 10 h dark cycle (white and grey background respectively). Traces represent mean ± SEM for larvae from the same clutch, genotype and treatment. n = 4 to 22 larvae per treatment and genotype. B: Log2FC between DMSO-treated WT and *srrm3*^ΔeMIC^ for the genes of significantly enriched cAMP pathway GO terms: “cyclic-nucleotide phosphodiesterase activity”, “cAMP metabolic process”. C: Maximum intensity z-projections of significant delta means obtained via MAPMapping^160^ of HCR *fosab*-stained WT and *srrm3*^ΔeMIC^ larvae at night shown dorsally (top) and sagittally (bottom). Sample size: N = 3 clutches with 6 to 10 larvae each per genotype. A, anterior; P, posterior; R, rightwards; L, leftwards; D, dorsal; V, ventral.

**Supplementary figure 7:**
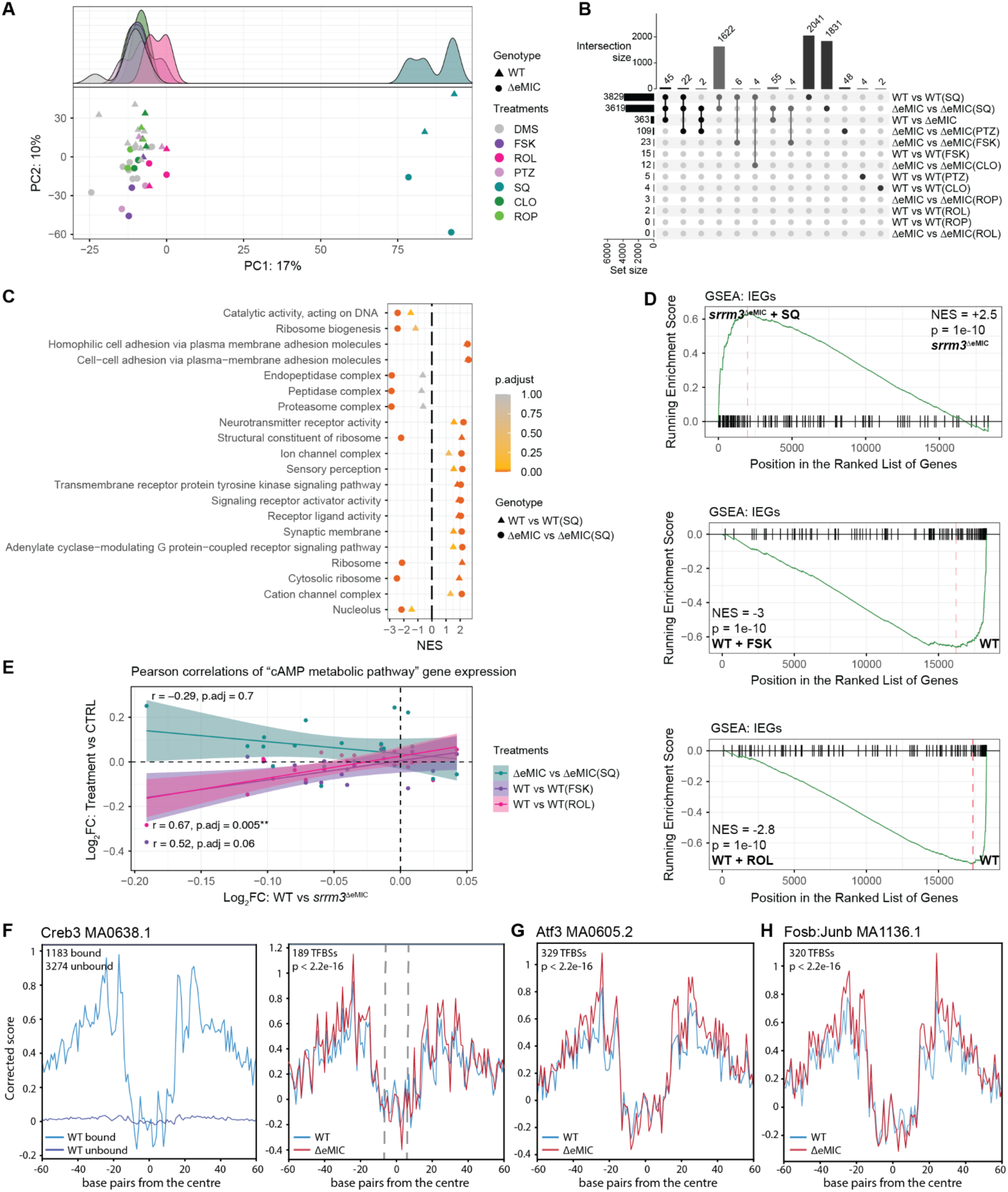
cAMP pathway modulators can partially mimic and reverse *srrm3*^ΔeMIC^- induced gene expression changes. A: PCA on the top 5000 most variable gene counts after variance stabilisation^152^ and batch correction with “Empirical Bayes-moderated adjustment for unwanted covariates”^153^. Density ridge plot shows PC1 separating SQ22536-treated and all other samples. Each dot represents one sample shaped by genotype and colored by treatment. B: Upset plot showing the overlap (min overlap = 2) of DEGs (p < 0.05) across conditions. Shades of grey indicate the number of overlapping sets (1 - 3). C: Top 20 GO terms (p.adj < 0.05) enriched among DEGs for ΔeMIC vs. ΔeMIC(SQ) based on NES scores obtained via GSEA (circles). Scores for WT vs. WT(SQ) are also indicated (triangles). Positive values indicate upregulation in the treatment condition. D: GSEA using “Mouse ortholog IEGs”^180^ as a test set, and using as background all expressed genes ranked by log2FC in gene expression between ΔeMIC vs. ΔeMIC(SQ) (top), WT vs. WT(FSK) (centre) and WT vs. WT(ROL) (bottom). E: Log2FC for “cAMP metabolic pathway” genes of the WT vs. *srrm3*^ΔeMIC^ comparison (x-axis) correlate with mimic groups [WT vs. WT(FSK): r = 0.52, p.adj = 0.06; WT vs. WT(ROL): r = 0.67, p.adj = 0.005] and anti-correlate with the rescue group [ΔeMIC vs. ΔeMIC(SQ): r = -0.29, p.adj = 0.7] (y-axis). Linear regression lines are shown. Statistics: Pearson correlations and p-values adjusted for testing across treatments using Bonferroni. F-H: Footprinting plots obtained from *TOBIAS*^176^. Genes associated with differential binding for any motif can be found in Table S3. N = 1 biological replicate with the number of transcription factor binding sites (TFBSs) indicated in the plots. F (left): showing all unbound and bound motifs of Creb3 binding sites in the WT. F (right): showing Creb3 binding sites bound stronger in the *srrm3*^ΔeMIC^ (log2FC > 0.25) compared to the WT. G: showing Atf3 binding sites bound stronger in the *srrm3*^ΔeMIC^ (log2FC > 0.25) compared to the WT. H: showing Fosb:Junb binding sites bound stronger in the *srrm3*^ΔeMIC^ (log2FC > 0.25) compared to the WT. P-values obtained using *TOBIAS* based on all TFBSs. A-E: Treatment abbreviations: CLO, clonidine; DMSO, dimethyl sulfoxide; FSK, forskolin; PTZ, pentylenetetrazol; ROL, rolipram; ROP, ropinirole; SQ, SQ22536.

